# The atypical antipsychotic aripiprazole alters the outcome of disseminated *Candida albicans* infections

**DOI:** 10.1101/2024.02.13.580133

**Authors:** Parker Reitler, Jessica Regan, Christian DeJarnette, Ashish Srivastava, Jen Carnahan, Katie M. Tucker, Bernd Meibohm, Brian M Peters, Glen E. Palmer

**Affiliations:** Integrated Program in Biomedical Sciences, College of Graduate Health Sciences, University of Tennessee Health Science Center, Memphis, Tennessee, USA; Pharmaceutical Sciences Program, College of Graduate Health Sciences, University of Tennessee Health Sciences Center, Memphis, Tennessee, USA; Department of Clinical Pharmacy and Translational Science, College of Pharmacy, University of Tennessee Health Sciences Center, Memphis, Tennessee, USA; Department of Pharmaceutical Sciences College of Pharmacy, University of Tennessee Health Sciences Center, Memphis, Tennessee, USA; Department of Microbiology, Immunology, and Biochemistry, College of Medicine, University of Tennessee Health Science Center, Memphis, Tennessee, USA

## Abstract

Invasive fungal infections (IFIs) impose an enormous clinical, social, and economic burden on humankind. For many IFIs, ≥ 30% of patients fail therapy with existing antifungal drugs, including the widely used azole class. We previously identified a collection of 13 approved medications that antagonize azole activity. While gain-of-function mutants resulting in antifungal resistance are often associated with reduced fitness and virulence, it is currently unknown how exposure to azole antagonistic drugs impact *C. albicans* physiology, fitness, or virulence. In this study, we examined how exposure to azole antagonists affected *C. albicans* phenotype and capacity to cause disease. We discovered that most of the azole antagonists had little impact on fungal growth, morphology, stress tolerance, or gene transcription. However, aripiprazole had a modest impact on *C. albicans* hyphal growth and increased cell wall chitin content. It also worsened the outcome of disseminated infections in mice at human equivalent concentrations. This effect was abrogated in immunosuppressed mice, indicating an additional impact of aripiprazole on host immunity. Collectively, these data provide proof-of-principle that unanticipated drug-fungus interactions have the potential to influence the incidence and outcomes of invasive fungal disease.

**Importance:** As natural inhabitants of the gastrointestinal and reproductive tracts, *Candida* sp. are routinely exposed to medications consumed by their human host. This study provides new insight into how drugs can modulate the physiology, fitness, and pathogenicity of *Candida albicans* - one of the most important human fungal pathogens. These results provide a proof of principle that co-administered medications consumed by at-risk patients may influence the initiation and/or outcome of life-threatening fungal disease.

## Introduction

Invasive fungal infections (IFIs) impose an enormous medical burden on humankind, causing an estimated 1.5 million deaths each year (1). *Candida albicans* is the most prevalent cause of IFIs in the western world, causing approximately 60,000 IFI’s annually in the United States alone, with associated costs estimated at approximately $2 billion (2–4). Three primary classes of antifungal drugs are used to treat IFI’s (5). The azoles inhibit synthesis of the membrane lipid ergosterol, the polyenes kill fungi through direct interactions with ergosterol that perturb membrane integrity (6), and the echinocandins disrupt cell wall synthesis through inhibition of β-1,3 glucan synthase (7). However, therapeutic failures occur with all three classes in ≥ 30% of the patients with disseminated *Candida* infections, and are even more common with other fungal pathogens (e.g *Aspergillus*, *Cryptococcus*). While acquired and intrinsic antifungal resistance may explain some cases of IFI treatment failure, most isolates remain sensitive, with just 0.5-2% of *C. albicans* isolates resistant to the azoles, and even less to the echinocandins (8–10). A variety of patient specific factors may contribute to poor outcomes, such as the severity of their immune dysfunction (11). Delays in diagnosis or in the provision of an appropriate antifungal agent is also associated with worsened outcomes (11). Yet many patients fail therapy for unknown reasons.

As eukaryotes, fungi and mammals share substantial similarity in their core metabolic and signaling modules (12–15). As such, it is likely that the physiology of fungi residing within the human body as part of the endogenous microbiota, or as invasive pathogens may be inadvertently targeted by the medications consumed by their host. Previously, we determined that a variety of drugs approved for human use can oppose the antifungal activity of fluconazole upon *C. albicans* (16) *in vitro*. Seven of these antagonists were selected for further evaluation in this study (Table 1), including the atypical antipsychotic aripiprazole. Our data reveal several antagonists elevated the minimum inhibitory concentration (MIC) of fluconazole by 8 - ≥ 256-fold. Further, the antagonistic activity of six of these drugs was dependent on the Tac1p zinc-cluster transcription factor, which activates expression of the ATP-driven Cdr1p and Cdr2p drug efflux pumps (17). Activity of the non-steroidal anti-inflammatory drug etofenamate was Upc2p-dependent, a transcription factor that activates expression of genes encoding enzymes required for ergosterol biosynthesis (18). Genetically encoded antimicrobial resistance - including azole-resistance in *C. albicans* conferred by activating mutations in the Tac1p and Upc2p transcription factors - is often associated with fitness costs (19,20). However, the impact of the fluconazole antagonistic drugs upon fungal physiology, pathogenicity, or fitness is unknown. The goal of this study was to investigate how fluconazole antagonistic medications affect the physiology, fitness, and pathogenicity of *C. albicans*, and determine if they can influence the cause of invasive infections, or the efficacy of antifungal therapy. Our next studies have sought to identify potentially unaccounted drug-fungus interactions and evaluate if the medications consumed by individual patients have the potential to influence the incidence or outcome of IFIs.

**Table 1.** Observed mass transitions and their monitoring parameters.

| Analytes | Precursor ion | Product ion | Declustering potential (V) | Collision energy (eV) |
| --- | --- | --- | --- | --- |
| **IS VERA** | 455.0 [M+H]<sup>+</sup> | 164.9 | 91 | 37.0 |
| **ARP1** | 449.3 [M+H]<sup>+</sup> | 393.1 | 60 | 24 |
| **ARP2** | 449.3 [M+H]<sup>+</sup> | 286.1 | 60 | 38 |
| **ARP3** | 449.3 [M+H]<sup>+</sup> | 219.2 | 60 | 36 |

**Table 1.** Medications that alter *Candida albicans* fluconazole sensitivity. *Increase in fluconazole MIC_S0_ of *C. albicans* strain SC5314 in the presence of 5 µM of the indicated drug, as determined using the CLSI antifungal susceptibility testing protocol 24-hours post incubation.

| Drug | Indication/class | Increase in MIC (Fold)* | Mechanism |
| --- | --- | --- | --- |
| Aripiprazole | Anti-psychotic | 32 | Tac1p-dependent |
| Penbutolol | β-blocker for blood pressure | 32 | Tac1p-dependent |
| Pinaverium | Anti-plasmodic | 32 | Tac1p-dependent |
| Mometasone | Anti-inflammatory glucocorticoid | 16 | Tac1p-dependent |
| Prenylamine | Vasodilator | 8 | Tac1p-dependent |
| Thiethylperazine | Anti-emetic | 8 | Tac1p-dependent |
| Etofenamate | Non-steroidal anti-inflammatory | >256 | Upc2p-dependent |

## Materials and Methods

### Growth conditions

*C. albicans* was routinely grown on YPD medium (1% yeast extract, 2% peptone, 2% dextrose) at 30°C, supplemented with uridine (50 μg/ml) when necessary. Transformant selection was carried out on minimal yeast nitrogen base (YNB) medium (6.75 g/liter yeast nitrogen base without amino acids, 2% dextrose, 2% Bacto agar), supplemented with the appropriate auxotrophic requirements as described for *Saccharomyces cerevisiae* (21) or 50 µg/ml uridine.

### Plasmid construction *Candida albicans* strain construction

The CAI4+pKE4+NRG1 strain was kindly provided by Brian M Peters. Plasmid pLUX (22) was also kindly provided by William Fonzi (Georgetown University). The *tac1*Δ/Δ and derived revertant strain were constructed using the auxotrophic marker system previously described by our laboratory using the TAC1DISF and TAC1DISR primers to delete the open reading frame, and TAC1AMPF and TAC1AMPR primers to select the open reading frame and 5’ promoter region for amplification into the pLUX plasmid (22). All oligonucleotides used in this study are listed in Table S8.

### Drug stocks

Stock solutions of fluconazole and each compound (aripiprazole, pinaverium, prenylamine, etofenamate, thiethylperazine, mometasone, and penbutolol) were prepared at 10 mM in dimethyl sulfoxide (DMSO) and diluted to required working concentrations.

### Growth kinetic assays

Growth curve assays were set up in 96-well plates using Roswell Park Memorial Institute (RPMI) buffered to pH 7 with (3-(N-morpholino)propanesulfonic acid). *C. albicans* strains were grown overnight in YPD at 30°C and the cell density was adjusted to 1 x 10^4^ cells/ml in the appropriate medium for the growth kinetic assays. An aliquot of cell suspension (100 µl) with an equal volume of twice the desired drug concentration of each antagonist. Cells were then incubated at 35°C inside a BioTek Cytation 5 plate reader shaking for 24-hours, and OD_600nm_ read every 30 min. Data was then analyzed via GraphPad Prism software. These assays were repeated in technical triplicate and in biological triplicate.

### Dry weight experiment

*C. albicans* strain SC5314 was grown overnight in YPD at 30°C. Cells were to inspect morphology. Remaining cells were then filtered with a 47 mm filter using a Whatman Swin-Lok filter set up, and lyophilized using a BenchTop Pro with Omnitronics before being weighed. Individual points indicate data from six independent experiments and cross bars indicates the mean.

### Phenotypic assays

*C. albicans* SC5314 was grown overnight in YPD at 30°C. The cell density of each and NaCl (up to 1 mM; ionic stress). Cells were plated, grown for 24-hours at 35°C, and scanned using biological duplicate.

### Agar hyphal growth assays

*C. albicans* SC5314 was grown overnight in YPD at 30°C before being washed in sterile deionized water. The cell density was adjusted to 1 x 10^7^ cells/ml in PBS and 2 µl of cell suspension spotted onto M199 (2% agar plates) supplemented with 5 µM of drug, or 0.5% DMSO (vehicle control). Plates were incubated for 96-hours at 37°C and imaged as above.

### RNA sequencing analysis

SC5314 was grown in YPD medium at 30°C overnight, then sub-cultured at purified from total RNA using poly-T oligo attached magnetic beads. After fragmentation, the first strand cDNA was synthesized using random hexamer primers, followed by the second strand cDNA synthesis using either dUTP for directional library or dTTP for non-directional library. Library was checked with Qubit and real-time PCR for quantification and bioanalyzer for size distribution detection. Libraries were then pooled and sequence on Illumina platforms. Genes were mapped to the SC5314 haploid genome value was converted to log_2_fold for the term “AVG log_2_fold”.

### Antifungal susceptibility testing

Antifungal susceptibility testing was performed using the broth microdilution method as described in CLSI document M27-A3 (24), with minor modifications. Fluconazole was diluted in DMSO, resuspended in RPMI-pH 7 at twice the final concentration, and serially diluted. *C. albicans* strains were grown overnight in YPD at 30°C, resuspended at 1 x 10^4^ cells/ml in RPMI-pH 7, and 100 µl transferred to wells of a round bottom 96-well plate containing and equal volume of diluted fluconazole solution. The final concentration of DMSO was 0.5% for all treatments, with drug-free control wells having DMSO alone. Plates were incubated without shaking for 24-hours at 35°C, and then scanned using an EPSON Perfection v700 Photo scanner. Experiments were performed in biological duplicate.

### Cell wall staining experiments

SC5314 cultures were diluted 1:100 into 5 ml YNB-pH 7 supplemented with either 5 µM aripiprazole or 0.5% DMSO (vehicle control). Cells were incubated at 30°C for 24-hours before harvesting at 2000 RPM for 5 minutes. Supernatant was removed and cells preserved in 10% formalin for 30 minutes on ice, washed three times, and stored in PBS. Cells were then counted and diluted to a final density of 1 x 10^6^ cells/ml in PBS. After centrifuging at 13,000 RPM for 1 minute, supernatant was removed and cells stained with either concanavalin A-FITC (50 µg/ml), wheat germ agglutinin-Alexa Fluor 488 (50 µg/ml), calcofluor white (50 µg/ml) or aniline blue (1 mg/ml) for 5 minutes, 1 hour, 30 minutes, or 1 hour respectively. For each stained sample, an unstained control was also prepared. After centrifugation, cells were resuspended into 500 µl PBS. Samples were then analyzed using a NovoCyte 3000 flower cytometer, using strain-specific lasers and filter sets. Mean fluorescence intensity (MFI) was compared between vehicle or drug-treated cells. Individual points indicate data from independent biological experiments and cross bars indicates the mean.

### Cultivation of cell lines

THP-1 macrophage-like cells (ATCC TIB-202) were cultured according to the manufacturer’s protocol in RPMI 1640 with 25 mM HEPES supplemented with 10% heat-inactivated fetal bovine serum (FBS), and 100 U/ml Penicillin-Streptomycin. (Pen-Strep) THP-1 cells were counted on Novocyte flow cytometer (Agilent) and frozen as aliquots of ∼5 x 10^6^ cells in liquid nitrogen.

### Macrophage challenge assays with live *C. albicans*

Upon recovery from cryopreservation, THP-1 cells were incubated for 3 days at 37°C, 5% CO_2_, and 90% humidity in complete RPMI 1640 medium containing 25 mM HEPES (10% heat-inactivated FBS and 100 U/ml Pen-Strep). After 3 days, THP-1 cells were counted on Novocyte flow cytometer (Agilent) and diluted to 2.5 x 10^5^ cells/ml in complete RPMI with 100 nM phorbol 12-myristate 13-acetate (PMA) (InvivoGen) to differentiate cells to a macrophage phenotype. Next, 200-μl aliquots were seeded at 5 x 10^4^ cells/well in 96-well tissue-culture treated polystyrene plates and incubated at 37°C with 5% CO_2_ for 24-hours. Overnight cultures of SC5314 were subcultured 1:100 into YNB-pH 7 and incubated at 30°C for 24-hours and washed 3 times in sterile PBS. *C. albicans* suspensions were then added to challenge medium containing 5 µM aripiprazole or 0.5% DMSO (vehicle control) in phenol red-free RPMI-pH 7 containing 25 mM HEPES to generate multiple multiplicities of infections (MOIs). Mock-infected controls using medium alone were also included. Drug-only controls were also included with THP-1 differentiated macrophages challenged with the same drug concentrations without SC5314. A positive control for TNF-α or IL-1β release was also prepared by challenging cells with 1 μg/ml lipopolysaccharide (*Escherichia coli* 0111:B4; InvivoGen) for an equivalent time, followed by addition of 5 mM ATP (InvivoGen) 30 min prior to the endpoint. The cells were challenged for 4-hours and gently centrifuged at 200 rpm for 2 minutes, and 100 μl of culture supernatant transferred to a polystyrene plate containing 100 µl of prediluted 1X ELISA/enzyme-linked immunosorbent spot (ELISPOT) assay buffer (eBioscience) and stored at −80°C. Culture supernatants were assessed for TNF-α or IL-1β using the Human Ready-Set-Go ELISA kit (eBioScience). ELISA optical density values from mock-infected controls were subtracted from those of *Candida*-challenged samples. Experiments were conducted in biological triplicate. Data are reported as the means + SEM.

### Macrophage challenge assays with fixed *C. albicans*

THP-1 cells were recovered, incubated, and differentiated with PMA as previously described above. For challenges with fixed SC5314 yeast cells, overnight cultures were subcultured at 1:100 into YNB-pH 7 and incubated with either 0.2, 1, 5 µM aripiprazole, or 0.5% DMSO (vehicle control) and incubated at 30°C for 24-hours. For challenges with fixed SC5314 germ-tubes, cells were subcultured at 1 x 10^7^ cells/ml in 50 ml RPMI-pH 7 (no phenol red) and incubated at 37°C for 1.5-hours. For both challenges, cultures were then centrifuged at 3000 RPM for 5 minutes, before supernatant was removed and cells preserved in 10% formalin on ice for 30 minutes. *C. albicans* was then centrifuged and washed 3 times in sterile PBS. Macrophages were challenged with an MOI of 20:1 with either 0.2, 1, 5 µM aripiprazole, or 0.5% DMSO (vehicle control) added to the challenge media as previously described. Similar controls were included as described above. Plates were either incubated for 24-hours (yeast) or 8-hours (germ-tube), supernatant collected, and ELISA assays for TNF-α and IL-1β performed as previously described. Experiments were conducted in biological triplicate. Data are reported as means + SEM.

### Ethics statement

The animals used in this study were housed in American Association for Accreditation of Laboratory Animal Care (AAALAC)-approved facilities at the University of Tennessee Health Science Center (UTHSC). The Institutional Animal Care and Use Committee (IACUC) at UTHSC approved the use of all animals and procedures (IACUC protocol number 22-0408). Mice were given standard rodent chow and water *ad libitum*. Mice were monitored daily for signs of distress, including noticeable weight loss and lethargy, and for the body condition score. The IACUC at UTHSC uses the Public Health Policy on Humane Care and Use of Laboratory Animals (PHS) and the Guide for the Care and Use of Laboratory Animals as a basis for establishing and maintaining an institutional program for activities involving animals. To ensure high standards for animal welfare, the IACUC at UTHSC remains compliant with all applicable provisions of the Animal Welfare Act (AWAR), guidance from the Office of Laboratory Animal Welfare (OLAW), and the American Veterinary Medical Association Guidelines on Euthanasia.

### Disseminate model of *C. albicans* infection

Mice were injected subcutaneously with 100 µl of either 10 mg/kg/day aripiprazole, 2 mg/kg/day aripiprazole, or vehicle twice daily, starting 3 days prior to infection. For drug formulations, aripiprazole (TCI America (98% Purity)) was prepared fresh daily as a formulation of 25% PEG400 (Sigma-Aldrich Cat. #. NC1211386), 19% Kolliphor RH40 (Sigma-Aldrich Cat. #. NC0542250) and 65% sterile water (Gibco Cat. #. 15230-147) and adjusted to 1 or 0.2 mg/ml, to achieve human equivalent steady state concentrations. The formulation was then sonicated and adjusted to appropriate volume of water before final sonication to obtain a clear solution. SC5314 was grown overnight in YPD broth at 30°C with shaking, washed twice in sterile PBS and resuspended at appropriate densities for infection of BALB/c and C57BL/6 mice (2.5 x 10^6^ cells/ml) or CD-1 mice (4 x 10^6^ cells/ml). Groups (n ≥ 8 mice) were inoculated by lateral tail vein injections with 100 µl of the desired cell suspension. Mice were monitored three times a day and those experiencing severe distress were humanely euthanized. For select experiments mice were euthanized at set time points of 48-hours post infection (p.i) for BALB/c and C57BL/6, or 72-hours p.i, for CD-1 strains. At endpoint, kidneys were extracted. Half of the left kidney was stored in 10% formalin for histopathology analysis, and remaining kidneys were homogenized in pre-weighed tubes of PBS. Serial dilutions of kidney homogenates were plated on YPD agar plates containing 50 μg/ml of chloramphenicol. Fungal burdens (CFU/g of kidney tissue) were determined by enumerating colonies formed after 48-hours of incubation at 30°C. For some experiments, mice were rendered leukopenic with 150 mg/kg cyclophosphamide in 200 µl of filter-sterilized PBS via intraperitoneal injections every 3 days, beginning 2 days prior to infection. Inoculum sizes were reduced to 2.5 x 10^5^ cells/ml for infections of BALB/c, or 4 x 10^5^ cells/ml CD-1 mice.

### Serum clinical chemistry

Blood was collected via cardiac puncture from each mouse at the time of sacrifice, and aliquoted into two vials (plasma and serum separation tubes-PST and SST) the blood in SST tubes was allowed to coagulate on ice for 30 minutes, and centrifuged at 1500 x g for 15 minutes. The blood in PST was centrifuged immediately at 3000 x g for 5 minutes and the plasma in supernatant was collected. Collected serum and plasma was stored at -80°C until ready for use. Serum samples were analyzed by UTHSC Regional Biocontainment Laboratory staff as fee-for-service using the DiaSys Diagnostic Systems chemical analyzer for blood urea nitrogen (BUN) and creatine.

### Quantification of aripiprazole concentration in plasma

Sample preparation was done by protein precipitation using ice-cold methanol containing verapamil (25 ng/ml) as internal standard. Each plasma sample (25 µl) was precipitated with 8-volumes (200 µl) of internal standard (IS), in methanol. Samples were vortexed for 30 seconds and filtered using 96-well Millipore sample filter plates, centrifuged at 4000 RPM for 5 min at 4°C and the filtrates were analyzed using LC-MS/MS. Chromatographic separations were carried out using a Shimadzu Nexera XR (LC-20ADXR) liquid chromatograph (Shimadzu Corporation, USA) consisting of two pumps, online degasser, system controller and an auto sampler. Mobile phase consisting of a) 95% water and 5% acetonitrile with 2 mM ammonium formate buffer and 0.1% formic acid and b) 95% acetonitrile and 5% water with 2 mM ammonium formate buffer and 0.1% formic acid was used at a flow rate of 0.6 mL/min in gradient mode. A Phenomenex® C18 2.6μ, 100 x 4.6 mm column (Phenomenex, USA) was used for separation. Samples (5 µl) were injected on column and the eluate was led directly into an API 4500 triple quadruple mass spectrometer (Applied Biosystems, Foster City, CA) equipped with a turbospray ion source was operated in the positive ion mode.

The concentration was determined based on the peak area ratio of IS vs analyte (verapamil vs aripiprazole) against the calibration curve. A suitable statistical regression model was chosen to after examination of residuals and coefficient of correlation. The lowest limit of quantitation (LLOQ) measured with acceptable accuracy and precision for aripiprazole from mouse plasma was established as 1.72 ng/ml. Appropriate dilution of samples was performed where necessary.

### Mouse tissue histology

Mice were euthanized 48-hours (p.i.), and half of the left kidney was cut cross-sectionally and stored in 10% formalin. Kidney samples were embedded in paraffin blocks and 5 µm slices prepared using a microtone. Serial sections were stained with hematoxylin and eosin (H&E) or Grocott-Gomori methenamine silver (GMS). Slides were scanned at 10X and 40X using a Nano Zoom-SQ Hamamatsu digital slide scanner.

## Results

### A subset of azole-antagonistic drugs impact *Candida albicans* morphogenesis

We initially determined whether exposure to the seven selected azole-antagonistic drugs (table 1) affected *C. albicans* growth. The SC5314 reference strain was suspended in RPMI medium with 5 µM of each drug or with 0.5% DMSO (vehicle control) and incubated at 35°C. Growth was then quantified at 30-minute intervals as OD_600nm_. Only aripiprazole and pinaverium had detectable effects on OD_600nm_, with both appearing to elevate *C. albicans* growth compared to drug-free control (fig. 1A). To further the impact of aripiprazole and pinaverium on *C. albicans* biomass, dry weights were determined after cells grown in RPMI with either drug or 0.5% DMSO (vehicle control) at 35°C. Neither drug increased dry mass compared to the vehicle control. In fact, aripiprazole slightly, but reproducibly, reduced dry mass (fig. 1B). The apparent contradiction between the drugs impact upon OD_600nm_ and biomass may be explained reflective of morphotype-specific effects given the capacity of *C. albicans* to switch between yeast and hyphal forms, which makes a major contribution to *C. albicans* virulence (25). To determine if any of the antagonists affect *C. albicans* morphogenesis, SC5314 was induced to form hyphae on M199 agar supplemented with 5 µM of each azole-antagonist, or 0.5% DMSO (vehicle control). After 96-hours at 37°C, a prominent border of hyphae could be observed at the margin of colonies formed in the absence of drug, as well as in the presence of five drugs. However, the hyphal growth was notably diminished in the presence of aripiprazole and pinaverium (fig. 1C). Similarly, aripiprazole and pinaverium suppressed hyphal growth in liquid RPMI medium (fig. 1D).

**Figure 1.**
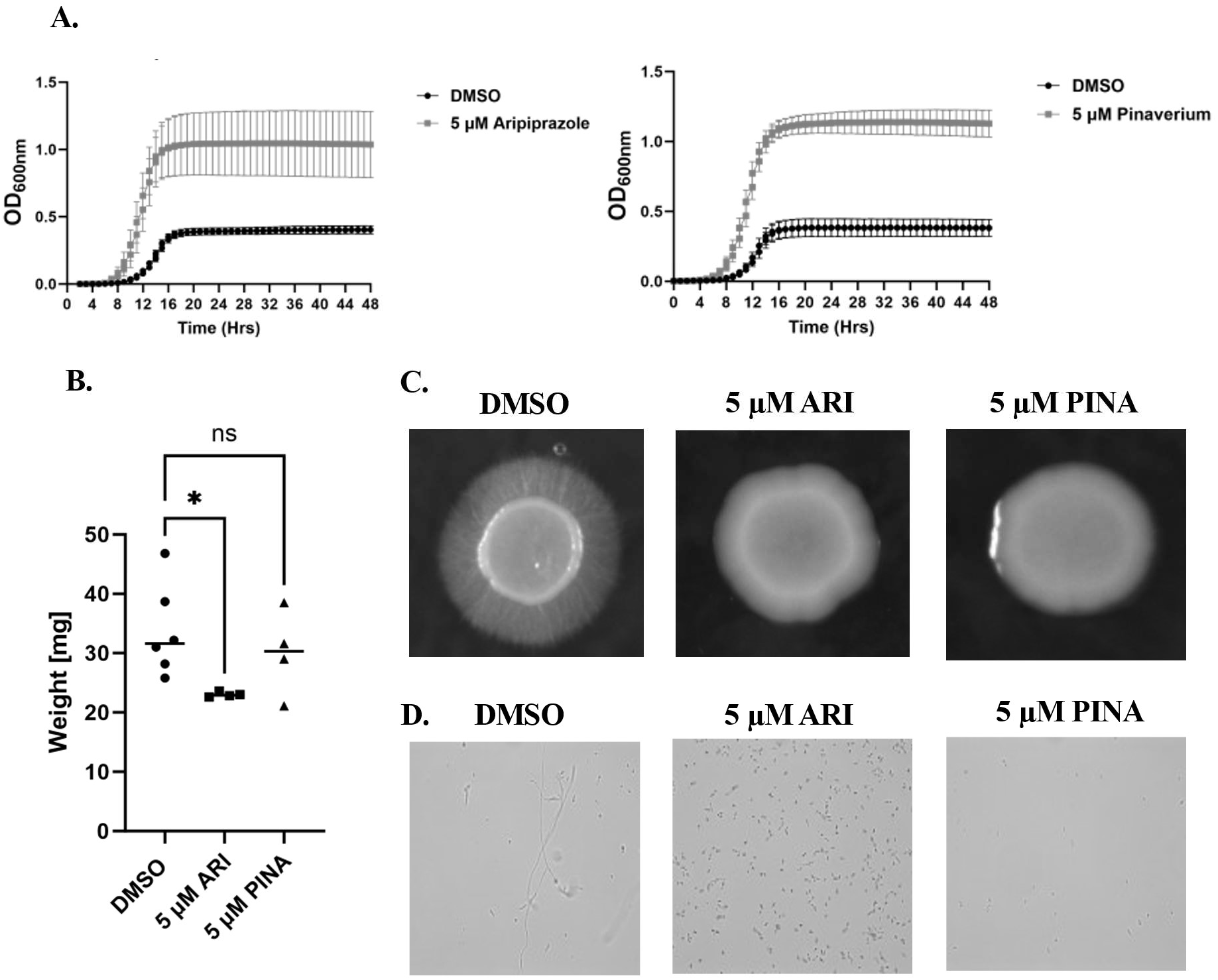
Aripiprazole and pinaverium alter *Candida albicans* cellular morphology. **A.** *C. albicans* strain SC5314 was subcultured at 1 x 10^4^ cells/ml in RPMI-pH 7 with 5 µM of aripiprazole (ARI), pinaverium (PINA), or 0.5% DMSO (vehicle), and incubated at 35°C for 48-hours. Growth was monitored as OD_600nm_ every 30 minutes. Experiments were performed in biological triplicate and plots indicate the mean + standard deviation for the OD_600nm_ value at each timepoint. **B.** SC5314 was subcultured at 1 x 10^6^ cells/ml in RPMI-pH 7 with 5 µM of either drug or vehicle and incubated at 35°C for 16-hours. Cells were harvested on filters, lyophilized, and weighed. Individual points indicate data from independent experiments and cross bar indicates the mean. Significance was calculated using a one-way ANOVA and Dunnet’s post-test **C.** SC5314 was suspended at 1 x 10^7^ cells/ml in PBS and 2 µl spotted on M199 agar supplemented with 5 µM of either drug or vehicle. Colonies were imaged after 96-hours incubation at 37°C. **D.** SC5314 was suspended in RPMI-pH 7 with either drug or vehicle and incubated at 35°C for 16-hours, fixed in 10% formalin, and cellular morphology examined microscopically with a 40X objective. ***\* P*** < 0.05. ns = not significant

We next asked whether exposure to the fluconazole antagonists affects *C. albicans* stress tolerance. SC5314 was grown in RPMI medium containing 5 µM of each antagonist or vehicle and supplemented with varying concentrations of NaCl (ionic stress), Congo red (cell wall stress), the detergent SDS (cell membrane stress), FeCl_3_ (electron-transport chain disruptor and oxidative stress) or caffeine (cell wall, cell membrane and DNA stressor). None of the seven drugs altered *C. albicans* sensitivity to NaCl, caffeine, FeCl_3_ or congo red (data not shown). However, treatment with all seven antagonists slightly increased sensitivity to SDS, with pinaverium having the greatest effect (fig. S1). Collectively, these data suggest that most of the antifungal antagonistic medications previously reported have little impact upon *C. albicans* growth, stress tolerance or morphogenesis. However, a subset including aripiprazole and pinaverium have a moderate impact on fungal physiology that is most notable during hyphal growth.

### Aripiprazole, mometasone and etofenamate regulate the transcription of a small and partially overlapping set of *Candida albicans* genes

To provide further insight into the effect of the azole-antagonists upon *C. albicans* physiology, we examined how aripiprazole, mometasone and etofenamate alter global patterns of gene transcription. SC5314 was grown at 35°C in RPMI medium supplemented with 5 µM of either drug, or 0.5% DMSO for 6-hours, total cellular RNA extracted and subject to high-throughput sequence analysis. The number of reads/kb were calculated for each gene and transcripts increased or decreased > 2-fold (*P* < 0.05) in the presence of drug (relative to the vehicle control) in each of 3 independently performed experiments considered responsive. Aripiprazole, mometasone, and etofenamate significantly increased the abundance of 31, 3 and 7 transcripts respectively (fig. 2A/2B). Notably, the abundance of 2 transcripts – those of *CDR2* and *RTA3* - were elevated by all three of the azole-antagonists (fig. 2C). *CDR2* encodes a multi-drug efflux pump of the ATP-binding cassette superfamily and has an established role in facilitating genetically encoded azole resistance in *C. albicans* (26), while *RTA3* encodes a phospholipid translocase also previously shown to affect azole-tolerance (27). These results are consistent with our previous data showing that the azole-antagonistic activity of aripiprazole is dependent upon the Tac1p transcription factor, which activates the transcription of *CDR2* (16). Several additional transcripts known to be either induced by Tac1p, or following exposure to fluphenazine (a known activator of Tac1p) (28), were identified as being elevated by one or more of the three drugs. These include the including that of *CDR1* transcript, encoding a second ATP-binding cassette type drug efflux pump that plays a major role in genetically encoded azole-resistance (26); and *PDR16,* that are responsive to both aripiprazole and etofenamate (table S1). Aripiprazole, mometasone and etofenamate decreased the abundance of 31, 22 and 10 transcripts respectively, with *HAK1* - encoding a putative potassium transporter - affected by all three (fig. 2C). Several, such as *MRV2*, have been reported to be either suppressed following fluphenazine exposure, by fluconazole exposure, or otherwise repressed in azole-resistant isolates (29). The relatively small sets of gene transcripts responsive to each of the azole-antagonists tested suggest a common mechanism underlying their activity. Further, the overlap in the transcripts regulated by more than one drug is especially surprising given their structural dissimilarity and diverse activity as both aripiprazole and mometasone are Tac1p-dependent, and etofenamate is Upc2p-dependent. Additionally, at least a subset of genes affected by one or more of the antagonists are associated with virulence. For example, *ICL1* (encoding isocitrate lyase) (30), *ALS1* (encoding a surface adhesin) (31), and *UME6* (encoding a transcription factor required to sustain hyphal growth) (32) transcripts are all suppressed by aripiprazole and have an established roles in supporting *C. albicans* virulence (table S4). Together, these data suggest that the drugs identified as undermining azole antifungal activity *in vitro* could potentially affect the outcome of invasive *C. albicans* infections through either modulating therapeutic efficacy, or the inherent virulence of the fungus itself. As we have previously shown that aripiprazole exerts azole-antagonistic activity within therapeutically relevant concentrations (i.e. that achieved within the bloodstream of patients on a standard dosing regimen: 150-450 ng/ml) we focused on identifying the underlying mechanism(s) (33).

**Figure 2.**
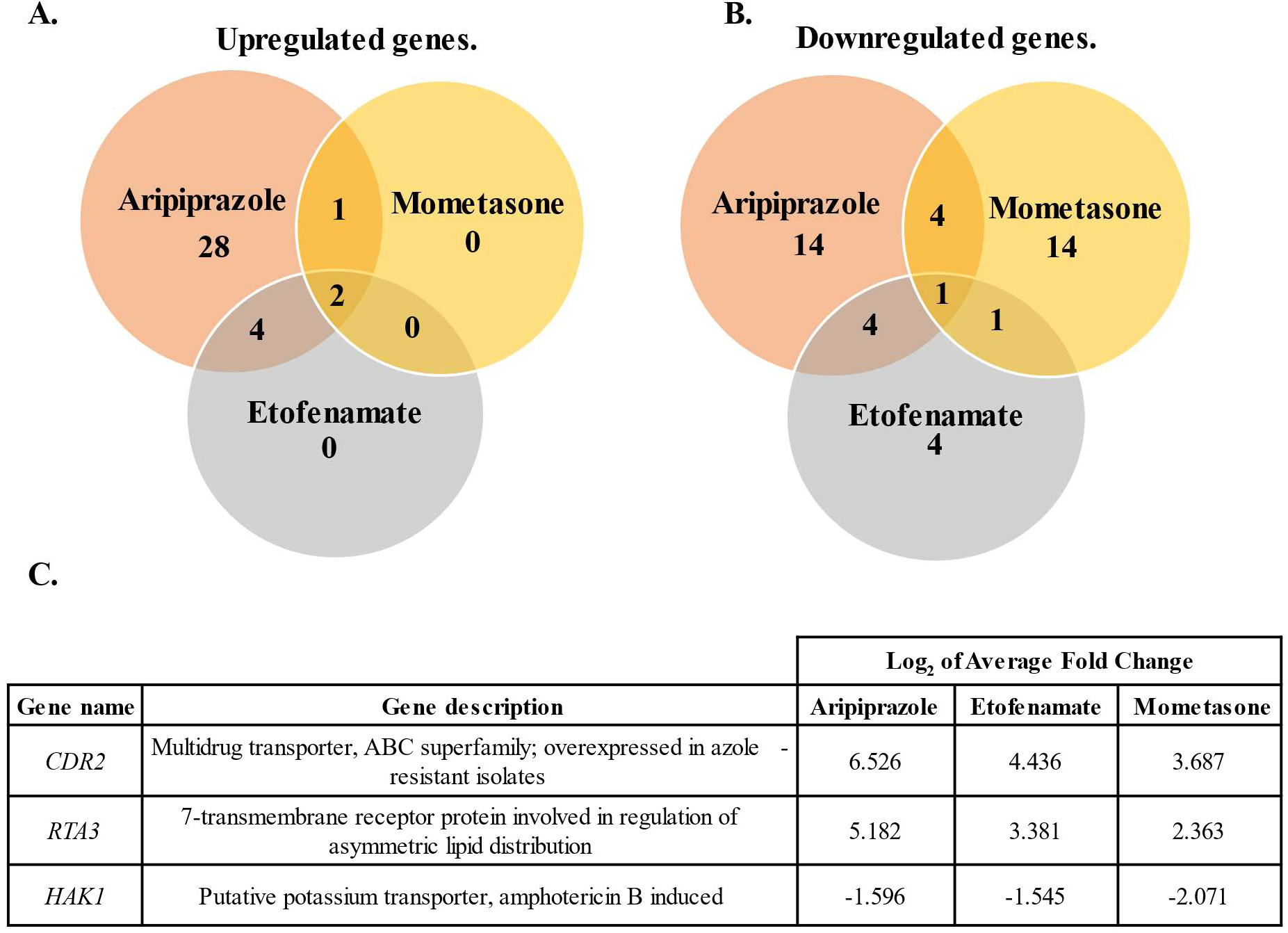
Azole-antagonistic drugs alter the transcription of a small and partially overlapping set of *Candida albicans* genes. SC5314 was subcultured into RPMI-pH 7 supplemented with 5 µM of aripiprazole, mometasone, etofenamate, or 0.5% DMSO (vehicle) and cultured for 6-hours at 35°C. Total RNA was extracted and transcriptional profiling performed. Drug responsive genes were defined as those increased or decreased by > 2-fold (adjusted p < 0.05) compared to vehicle treated samples in each of 3 independent biological replicates **A-B.** Venn diagram showing the number of gene transcripts for significantly upregulated genes **(A.)** or downregulated genes **(B.) C.** Genes identified that were responsive to all three drugs.

### Aripiprazole-related drugs oppose fluconazole antifungal activity through two distinct mechanisms

Aripiprazole is composed of 3,4-Dihydro-2(1H)-quinolinone and (2,3-Dichlorophenyl)piperazine moieties coupled by a butoxy-linker (fig. 3A). To determine which chemical structure was responsible for its Tac1p-dependent azole-antagonistic activity, we evaluated the azole-antagonistic activity of two additional closely related drugs. Bexpiprazole has a benothiophene in place of the (2,3-Dichlorphenyl)piperazine moiety of aripiprazole and elevated the fluconazole MIC of *C. albicans* by ∼8-fold (fig. 3B). Cariprazine possesses the (2,3-Dichlorophenyl)piperazine structure but has an ethyl pinker and cyclohexyl and dimethylurea groups in place of the 3,4-Dihydro-2(1H)-quinolinone of aripiprazole (fig. 3A). Cariprazine also had marked fluconazole antagonistic activity on *C. albicans*, elevating the MIC by >256-fold (fig. 3B). These data suggest that the phenylpiperazine group common to all three drugs is likely responsible for the antagonistic activity. However, this interpretation may be oversimplified as (2,3-Dichlorophenyl)piperazine alone did not significantly affect *C. albicans* fluconazole sensitivity (fig. S3B). The antagonistic activity of aripiprazole is Tac1p-dependent, while Bexpiprazole was both Tac1p and Upc2p-dependent, and cariprazine’s entirely Upc2p-dependent (fig. 3B). These data are consistent with the 2-quinolinone group of aripiprazole and bexpiprazole activating Tac1p. However, neither the 3,4-Dihydro-2(1H)-quinolinone found in bexpiprazole, nor the 3-cyclohexyl-1,1-dimethylurea group found in cariprazine are sufficient to decrease fluconazole sensitivity (fig. S3C/S3E). Collectively these data underscore the fact that biologically active small molecules can affect *C. albicans* physiology and antifungal sensitivity through multiple mechanisms and that the structure-activity relationships underlying these interactions are complex and multifaceted.

**Figure 3.**
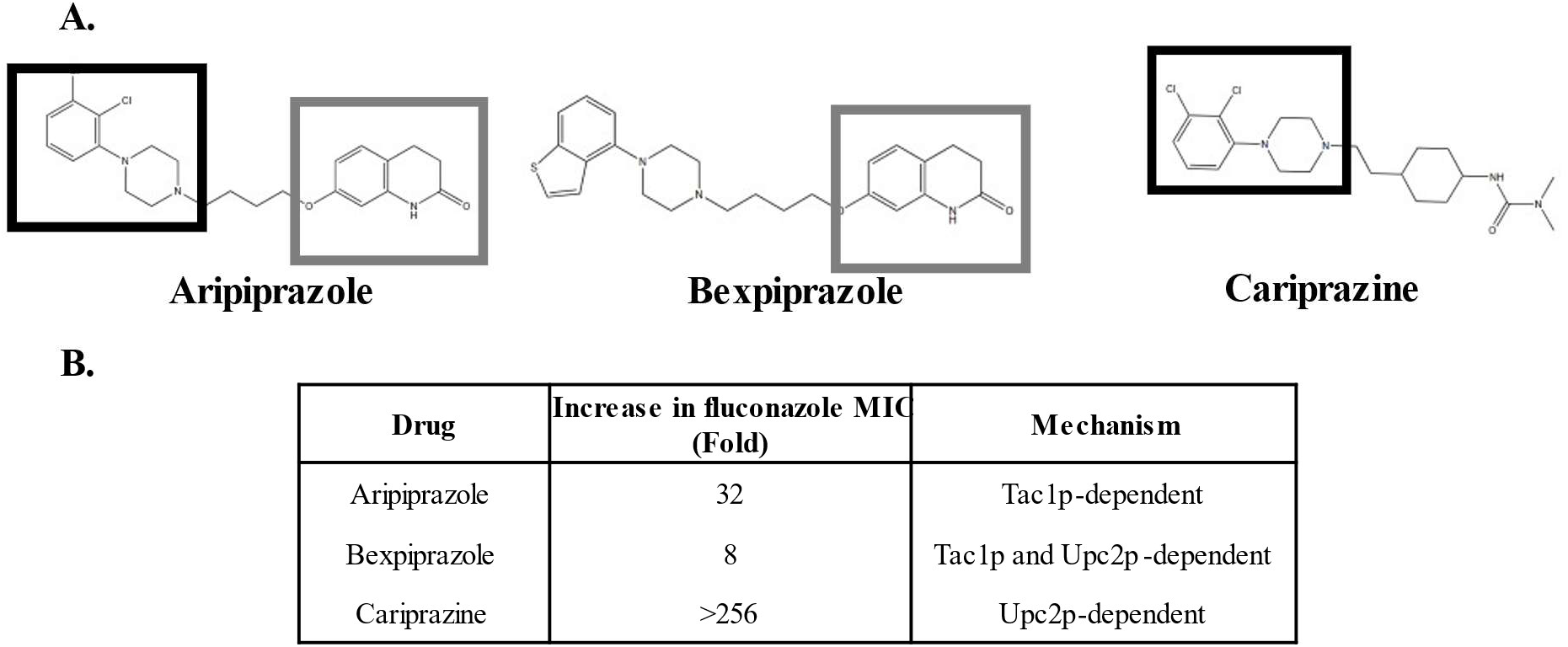
Aripiprazole related compounds also possess azole antagonistic activity. **A.** Structures of aripiprazole and structurally related drugs, bexpiprazole and cariprazine. Shared sub-structures are designated as follows: (2,3-Dichlorophenyl)piperazine (group indicated by black square), and 3,4-Dihydro-2(lH)-quinoline (indicated by gray square). **B.** Fluconazole antagonistic activity was determined by CLSI susceptibility assays conducted with *C. albicans* SC5314 as well as strains lacking either Taclp or Upc2p in RPMI-pH 7 with 5 µM of each compound or 0.5% DMSO (vehicle control). Minimum inhibitory concentrations (MICs) were denoted at 24-hours for each compound relative to the vehicle control. Experiments were run in biological duplicate.

### Aripiprazole does not affect the efficacy of fluconazole therapy in a mouse model of disseminated candidiasis

To determine if the antagonistic activity of aripiprazole observed *in vitro* affects the therapeutic efficacy of fluconazole during *in vivo* infection, we utilized a mouse model of disseminated candidiasis. Two groups of BALB/c mice were treated with either aripiprazole (5 mg/kg/day) or vehicle by subcutaneous injection for 3 days prior to infection with ∼ 2.5 x 10^5^ colony forming units (CFU) of SC5314 via the tail vein. Daily treatments with aripiprazole or vehicle were continued, and 24-hours post-infection each group was sub-divided - one sub-group was treated with fluconazole (5 mg/kg/day); and the second mock-treated with vehicle alone via intraperitoneal injection. Mice were euthanized 5-days post-infection (p.i), kidneys extracted, homogenized, and tissue fungal burden quantified as CFU. Fungal colonization was similar between the fluconazole and fluconazole + aripiprazole treated groups (fig. 4A), suggesting that aripiprazole does not antagonize fluconazole efficacy in this model. However, two mice in the aripiprazole treated group unexpectedly succumbed to infection prior to the scheduled endpoint of the experiment (data not shown). Strikingly, the remaining mice in this group displayed advanced signs of morbidity. Importantly, uninfected mice treated with aripiprazole exhibited no weight loss or other signs of disease (data not shown). While no *in vivo* azole antagonism was noted, these results unexpectedly demonstrated that aripiprazole may impact *C. albicans* virulence or the progression of invasive disease in the absence of azole-therapy.

**Figure 4.**
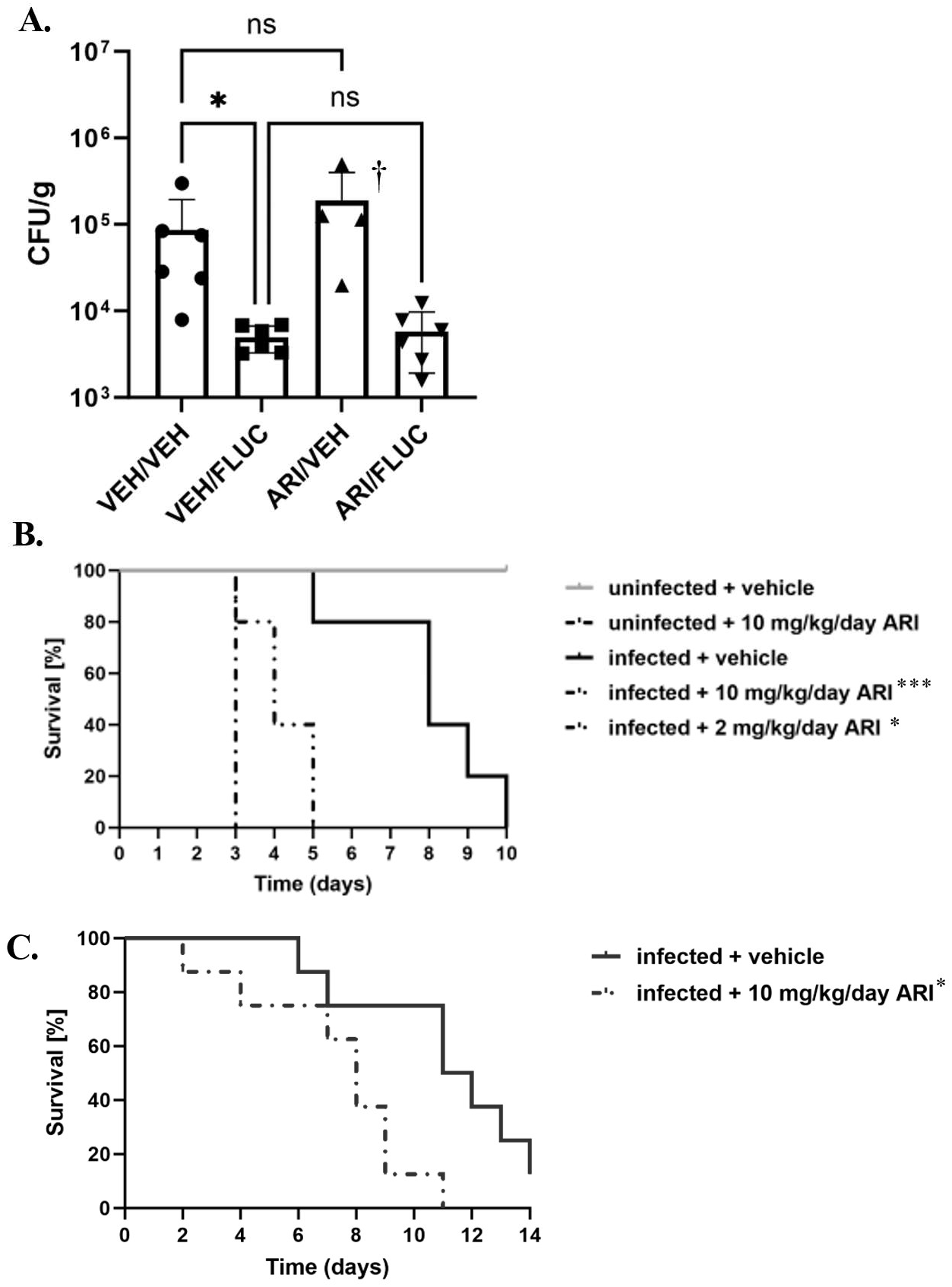
Aripiprazole exposure exacerbates the severity of disseminated *Candida albicans* infection in mice. **A.** 0, mFemale BALB/c mice (n = 6 per group) were treated daily with 5 mg/kg/day aripiprazole (ARI) or vehicle (VEH) starting 3 days prior to infection. Mice were then infected intravenously with ∼ 2.5 x 10^5^ of SC5314. Fluconazole (5 mg/kg/day - FLUC) was administered 24-hours post infection (p.i), and mice were euthanized five days p.i. Kidneys were extracted, homogenized, and fungal burden quantified as colony forming units (CFU) per gram of tissue. Data is depicted as the mean + standard deviation, and statistical significance was calculated using a one-way ANOVA and Kruskal-Wallis post-test. †Only four mice plotted for ARI/VEH group as succumbed prior to the endpoint. **B.** Groups of BALB/c mice (n > 11) were treated daily with 10 or 2 mg/kg/day aripiprazole (ARI) or vehicle starting 3 days prior to infection, and infected with 2.5 x 10^5^ of SC5314 via lateral tail vein injection and monitored for 10 days. The survival of the infected drug treated groups were compared to the infected mock treated group, using a log-rank test. **C.** Groups of CD-I mice (n = 8) were heated and infected as described in **(B.)** and survival monitored for 14 days. Survival was compared using a log-rank test. *P<0.05, *** ***P<*** 0.0001.

### Aripiprazole exposure exacerbates the severity of disseminated *Candida albicans* infection in mice

To further examine the effect of aripiprazole on the outcome of disseminated infection, groups of BALB/c mice were treated with either 2 or 10 mg/kg/day aripiprazole or vehicle alone, and after 3 days infected with SC5314 as described before. The mice were monitored daily for signs of disease progression, those exhibiting signs of distress euthanized, and survival compared using the log-rank test. Mice in both aripiprazole treated groups succumbed to infection more rapidly than the vehicle treated mice (fig. 4B), while uninfected mice treated with 10 mg/kg/day of aripiprazole again remained healthy throughout the experiment. Similar results were obtained using outbred CD-1 mice (fig. 4C), indicating these results are not mouse strain-dependent. Endpoint experiments at 48-hours p.i, kidneys revealed no significant differences in fungal burden between the three treatment groups in levels of fungal colonization measured as CFUs (fig. 5B). However, foci of fungal invasion the kidneys of both aripiprazole treated groups compared to mock treated animals (fig. 5A). Additionally, the kidneys of infected aripiprazole-treated mice demonstrated substantially larger inflammatory lesions characterized by extensive PMN infiltration than were observed in the mock treated group (fig. 5A). Analysis of serum samples obtained at 48-hours p.i. also revealed higher levels of blood urea nitrogen (BUN) than the mock treated control mice (fig. 5C/5D), providing additional evidence that aripiprazole exposure exacerbates kidney damage and dysfunction during disseminated candidiasis. Notably, aripiprazole treatment had no effect on BUN or creatine levels in uninfected mice. Comparable results were obtained with C57BL/6 mice, further supporting that the effect of aripiprazole on the outcome of systemic *C. albicans* infection is mouse strain-independent (fig. S4).

**Figure 5.**
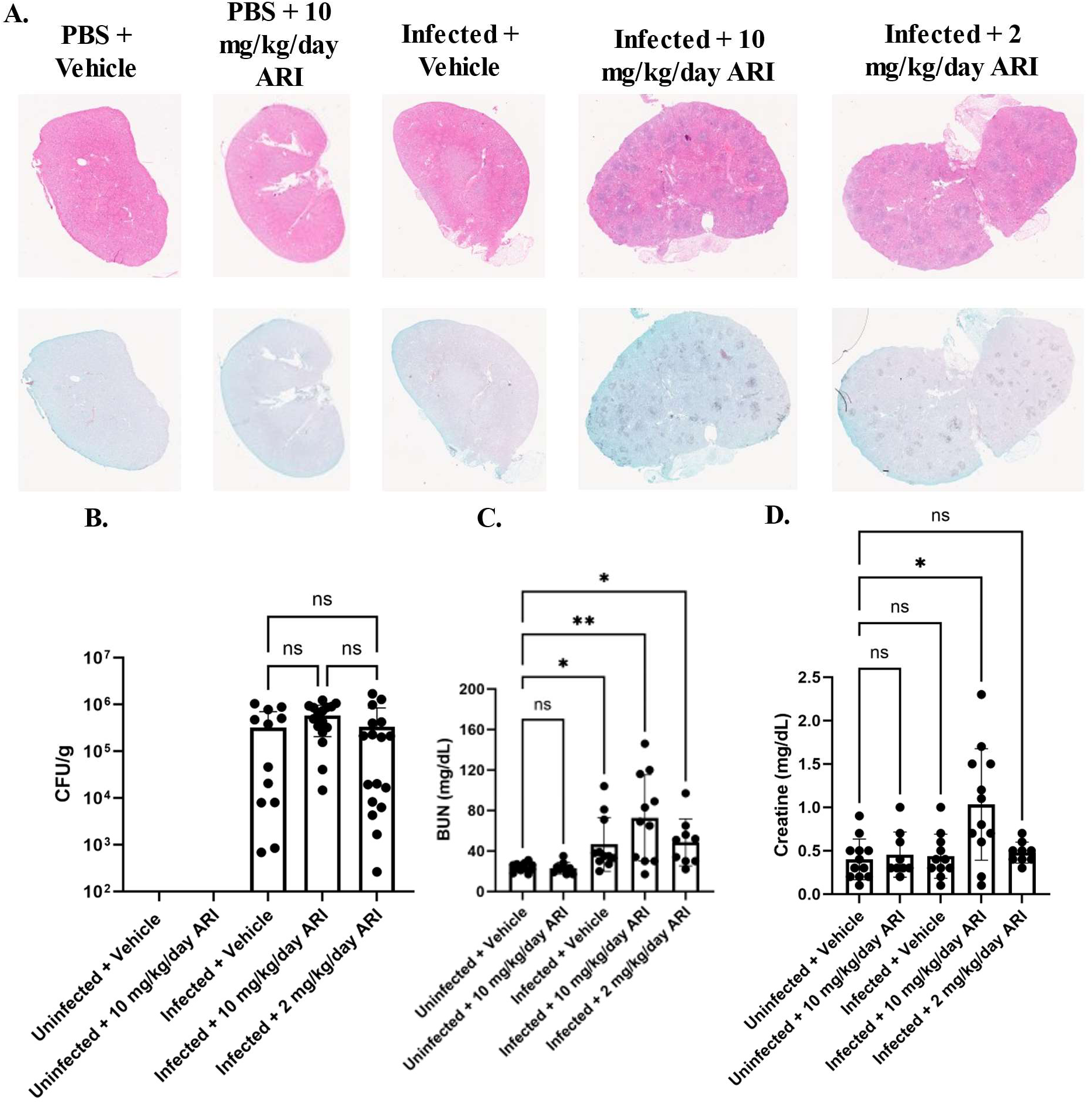
Aripiprazole treatment increases fungal invasion and kidney damage. **A.** BALB/c mice were treated daily with either 10 mg/kg/day or 2 mg/kg/day aripiprazole, or vehicle starting 3 days prior to infection. Mice were then infected with 2.5 x 10^5^ of SC5314 or PBS (uninfected control). At the 48-hour endpoint, mice were euthanized, kidneys extracted, and stained with H&E (top panel) or GMS (bottom panel) for each condition. Pictures are representative (n = 3 per group). **B.** Kidneys were extracted from the same mice and fungal burden was quantified as colony forming units (CFU) per gram of tissue. Data is depicted as the mean + standard deviation, and statistical significance calculated using a one-way ANOVA and Kruskal-Wallis post-test (n> 11). **C-D.** Blood was extracted from mice in **(A.).** Serum was processed, and blood urea nitrogen (BUN) **(C.)** or creatinine **(D.)** levels quantified. Groups were compared to uninfected vehicle. Data is depicted as the mean + standard deviation and significance determined using a one-way ANOVA and Kruskal-Wallis post-test (n >11). ***\* P*** < 0.05, ** ***P*** < 0.005. ns = not significant

Finally, we next examined how aripiprazole exposure affects disease progression in severely immunosuppressed mice. Survival experiments were performed as described above, with BALB/c mice were rendered leukopenic through treatment with cyclophosphamide two days prior to infection and infected with 2.5 x 10^4^ CFU of SC5314 three days thereafter. Strikingly, the previously observed difference in survival between vehicle and aripiprazole treated groups (fig. 4B) was eliminated (fig. 6A). Similar findings were observed using outbred CD-1 (fig. 6B). In a separate 48-hour endpoint experiment, kidney fungal burden and histopathology were assessed in cyclophosphamide treated CD-1 mice. Interestingly, while pockets of fungal growth were observed throughout the kidneys in both aripiprazole and mock treated groups, there were no obvious zones of inflammation (fig. 6C). Quantitative fungal burdens supported the microscopy findings (fig. 6D). Additionally, BUN and creatine levels measured in serum samples taken 48-hours p.i were similar between aripiprazole and mock treated groups (fig. 6E). These data suggest that the impact of aripiprazole during systemic *C. albicans* infection is at least partially dependent on host immune status.

**Figure 6.**
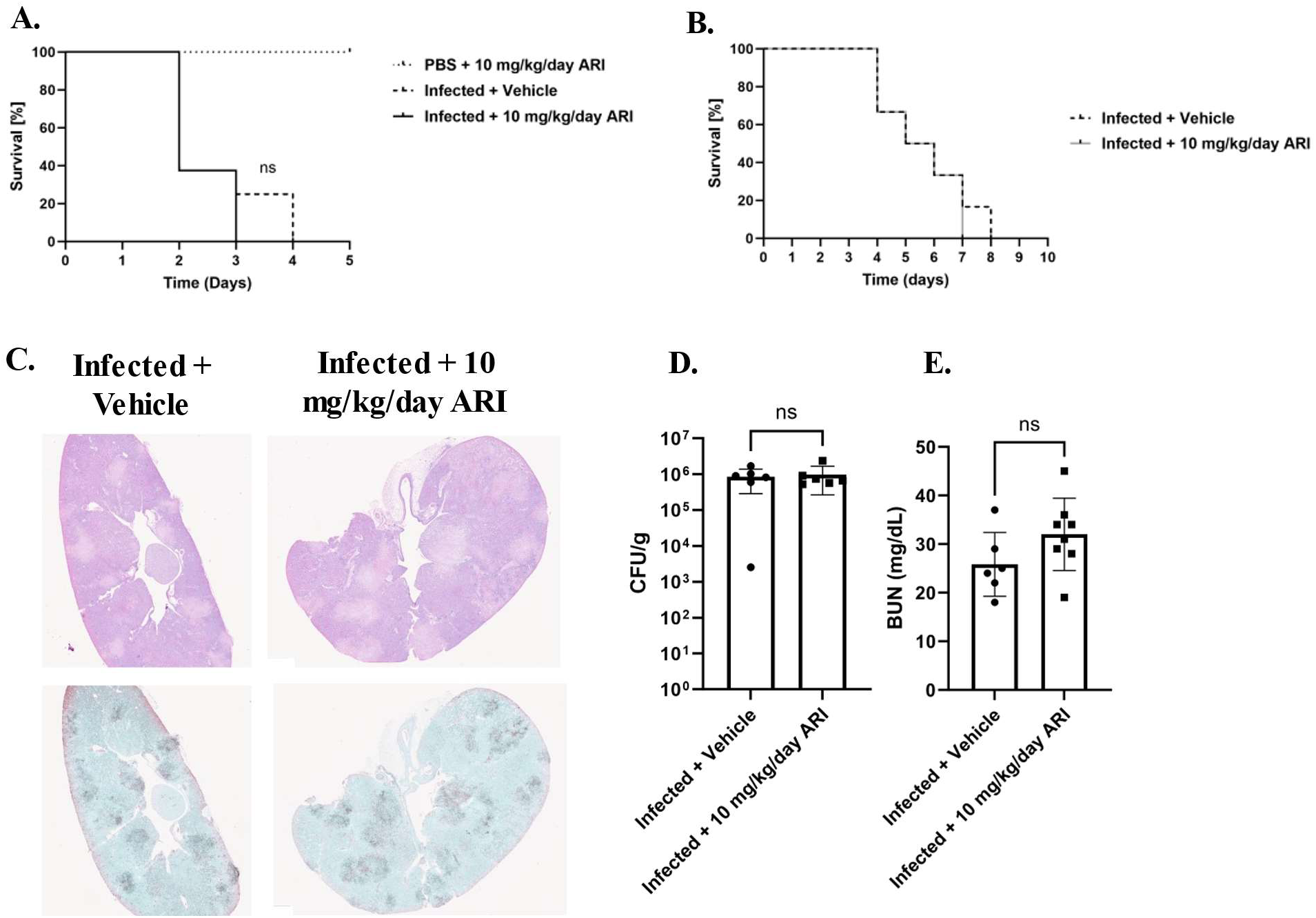
Aripiprazole does not affect outcome of disseminated *Candida albicans* infection in cyclophosphamide treated mice. **A.** BALB/c mice (n = 8 per group) were treated with either 10 mg/kg/day aripiprazole (ARI) or vehicle daily starting 3 days prior to infection. They were also administered cyclophosphamide (150 mg/kg) two days prior to infection and three days thereafter. Mice were then infected with 2.5 x 10^4^ of *C. albicans* SC5314 and monitored. The survival of the infected drug treated groups were compared to the infected mock treated group, using the log-rank test. **B.** Groups of CD-I mice (n = 8) were treated as described in **(A.),** infected with 5 x 10^4^of *C. albicans* SC5314, and survival monitored for 10 days. Survival was compared using a log-rank test. **C.** CD-I mice were treated and infected as described in **(B.)** and mice euthanized 72-hours p.i. Kidneys were extracted and stained with either H&E (top panel) or GMS (bottom panel). Pictures are representative (n = 3 per group). **D.** Kidneys were obtained from mice in **(C.)** and fungal burden quantified as CFU per gram of tissue. Data is depicted as the mean + standard deviation, and statistical significance calculated using a one-way ANOVA and Mann-Whitney post-test (n = 8). **E.** Blood was extracted from the same mice as in **(C.)** serum processed, and blood urea nitrogen (BUN) levels quantified. Groups were compared to infected treated versus 10 mg/kg/day aripiprazole infected mice. Data is depicted as the mean + standard deviation, and statistical significance was calculated using a one-way ANOVA and Mann-Whitney post-test (n = 8). ns = not significant

### Aripiprazole modulates host response to *Candida albicans*

Given that aripiprazole worsens infection outcome in an immune-dependent manner, we asked whether this drug alters immunogenicity of *C. albicans* by modulating cell wall pathogen molecular patterns (PAMPs). To determine this, *C. albicans* was grown in the presence or absence of 5 µM aripiprazole, fixed, and stained with aniline blue (total β-glucan), concanavalin A (total mannan), wheat-germ agglutinin (exposed chitin) or calcofluor white (total chitin), and relative fluorescence levels quantified by flow cytometry (fig.7). Aripiprazole had no obvious effect on β-glucan or mannan content, but did induce a significant increase in cell wall chitin content (∼1.5-fold versus vehicle treated control). We speculate that changes in polysaccharide content could modulate PAMP presentation at the *C. albicans* cell surface, and in turn the capacity of the mammalian host to detect or respond to the fungus. To test this hypothesis, we used an *in vitro* model of *C. albicans*-macrophage interaction. THP-1 derived macrophages were challenged with *C. albicans* at multiplicity of infections (MOIs) of either 1:1, 2:1, 4:1, or 8:1 in the presence of either 5 µM aripiprazole or DMSO (vehicle control). After 4-hours incubation, release of the pro-inflammatory IL-1β and TNF-α cytokines into the culture supernatant was compared by ELISA. Significantly less IL-1β (fig. 8A) and TNF-α (fig. 8B) were released in the presence of aripiprazole at MOIs of 2:1 or greater, indicating an altered host-fungus interaction. To determine if aripiprazole-mediated cytokine suppression was driven by its effect on host or fungal physiology, additional macrophage challenges were conducted using fixed *C. albicans*. Firstly, *C. albicans* SC5314 was grown as yeast with 0.2, 1 or 5 µM aripiprazole or vehicle (DMSO), fixed, washed, and then applied to macrophages. Fixed yeast failed to induce robust release of either TNF-α or IL-1β (fig. 9A). However, when *C. albicans* was as germ-tubes and subsequently fixed, those cultured in the presence of aripiprazole induced significantly less TNF-α or IL-1β than vehicle treated cells, with the differences most obvious at the highest concentrations (fig. 9B). This correlated with suppression of hyphal growth in the aripiprazole treated cells as described previously (fig. 1D). To examine if aripiprazole also impacts host response to *C. albicans*, untreated fungi were grown as previously described, fixed, washed, then applied to macrophages in the presence of 0.2, 1 or 5 µM aripiprazole or vehicle. Surprisingly, macrophages co-incubated with fixed germ-tubes in the presence of 5 µM aripiprazole, also released significantly less TNF-α, with a similar trend observed for IL-1β that did not reach statistical significance (fig. 9D). In contrast, macrophages co-incubated with fixed yeast in the presence of 5 µM of aripiprazole significantly increased release of both TNF-α and IL-1β (fig. 9C). Together, these results suggest that aripiprazole likely alters both host and fungal physiology during disseminated candidiasis to drive worsened outcomes.

**Figure 7.**
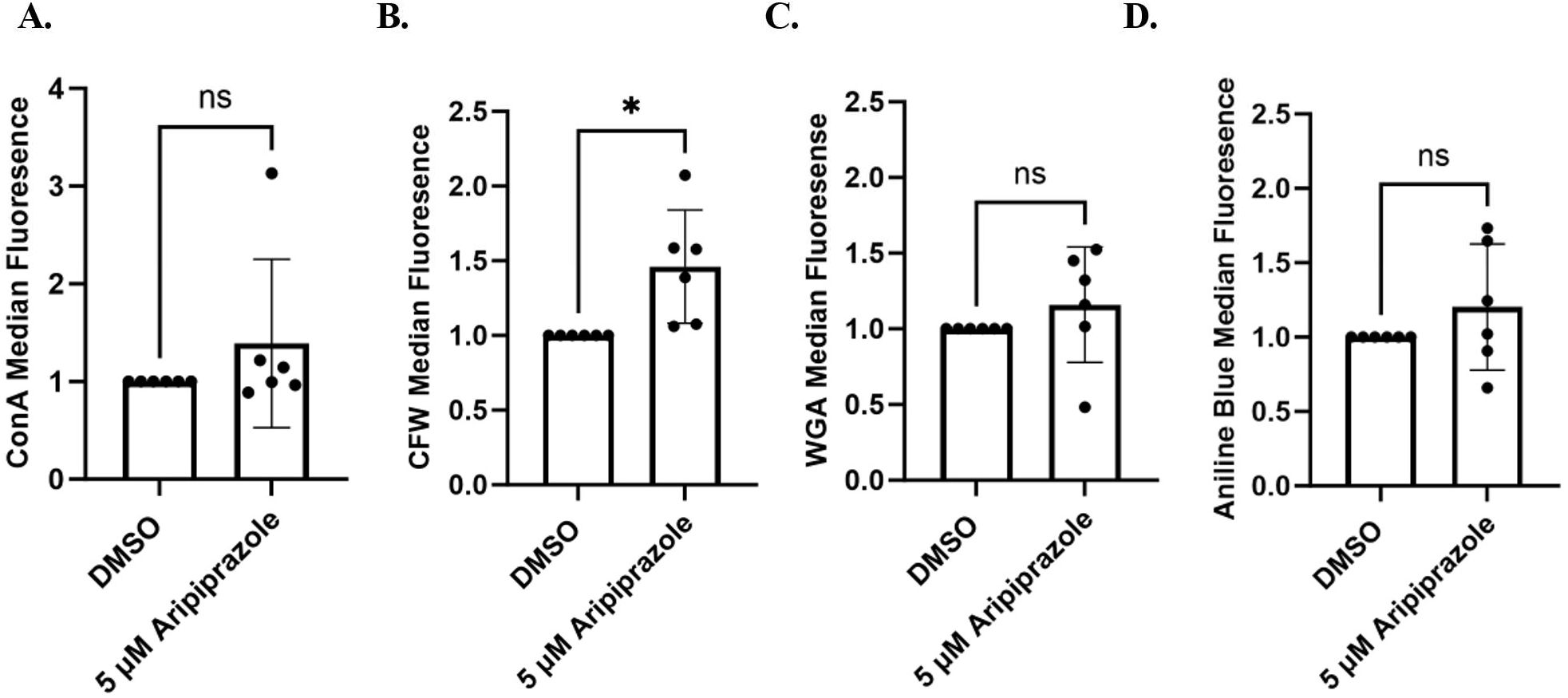
Aripiprazole modulates *Candida albicans* cell wall chitin content. SC5314 was cultured in YNB-pH 7 with either 5 µM of aripiprazole or 0.5% DMSO (vehicle) at 3O°C for 24-hours. Cells were then fixed in formalin, washed and stained for mannan (concanavalin A-FITC conjugate) **(A.),** chitin content (calcofluor white) **(B.),** chitin exposure (wheat germ agglutinin-Alexa Fluor 488 conjugate) **(C.),** or β-glucan content (aniline blue) **(D.).** Fluorescence intensity of was measured by flow cytometry and drug-treated groups normalized to the paired vehicle control. Each data point represents the median fluorescence of 20,000 events and data depicted as the mean + standard deviation of 6 independent experiments. Statistical significance was calculated using two-tail unpaired Student’s t-test. ***\* P <*** 0.05. ns = not significant

**Figure 8.**
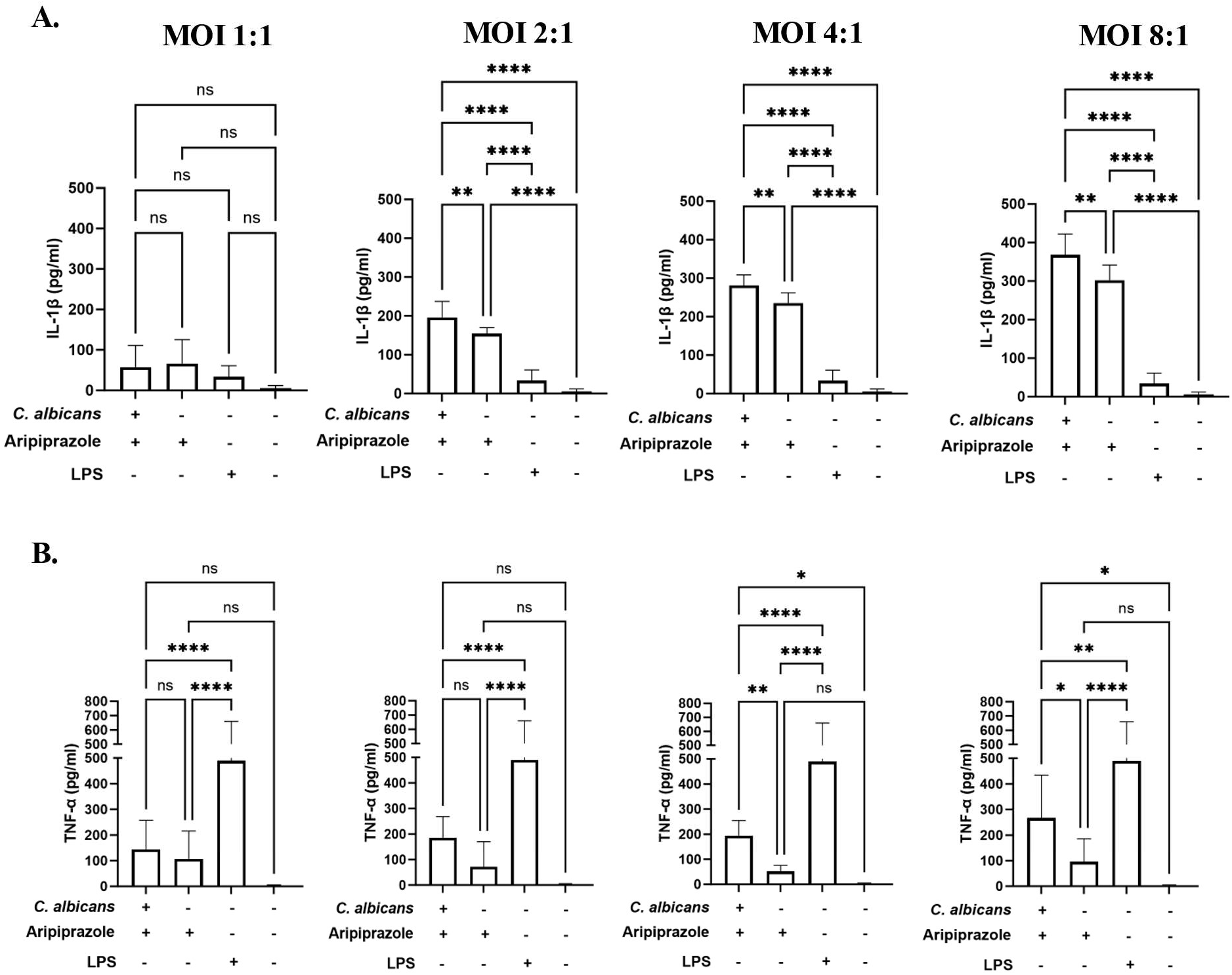
Aripiprazole decreases pro-inflammatory cytokine release during *Candida albicans-* macrophage interaction. **A.** Human THP-1 derived macrophages were co-incubated with SC5314 at the indicated MOI in RPMI-pH 7 medium supplemented with either 5 µM of aripiprazole or 0.5% DMSO (vehicle). Supernatants were collected after 4-hours co-incubation at 37^¤^C. ELISA assays were performed to quantify ΓL-lβ **(A.)** or TNF-α **(B.).** Experiments were conducted in technical quadruplicate and performed independently in biological triplicate. Data is depicted as the mean + standard deviation. Statistical significance was calculated using one-way ANOVA and Dunnet’s post-test. ***\*\* P <*** 0.005, *** ***P <*** 0.0005, **** ***P <*** 0.0001. ns = not significant

**Figure 9.**
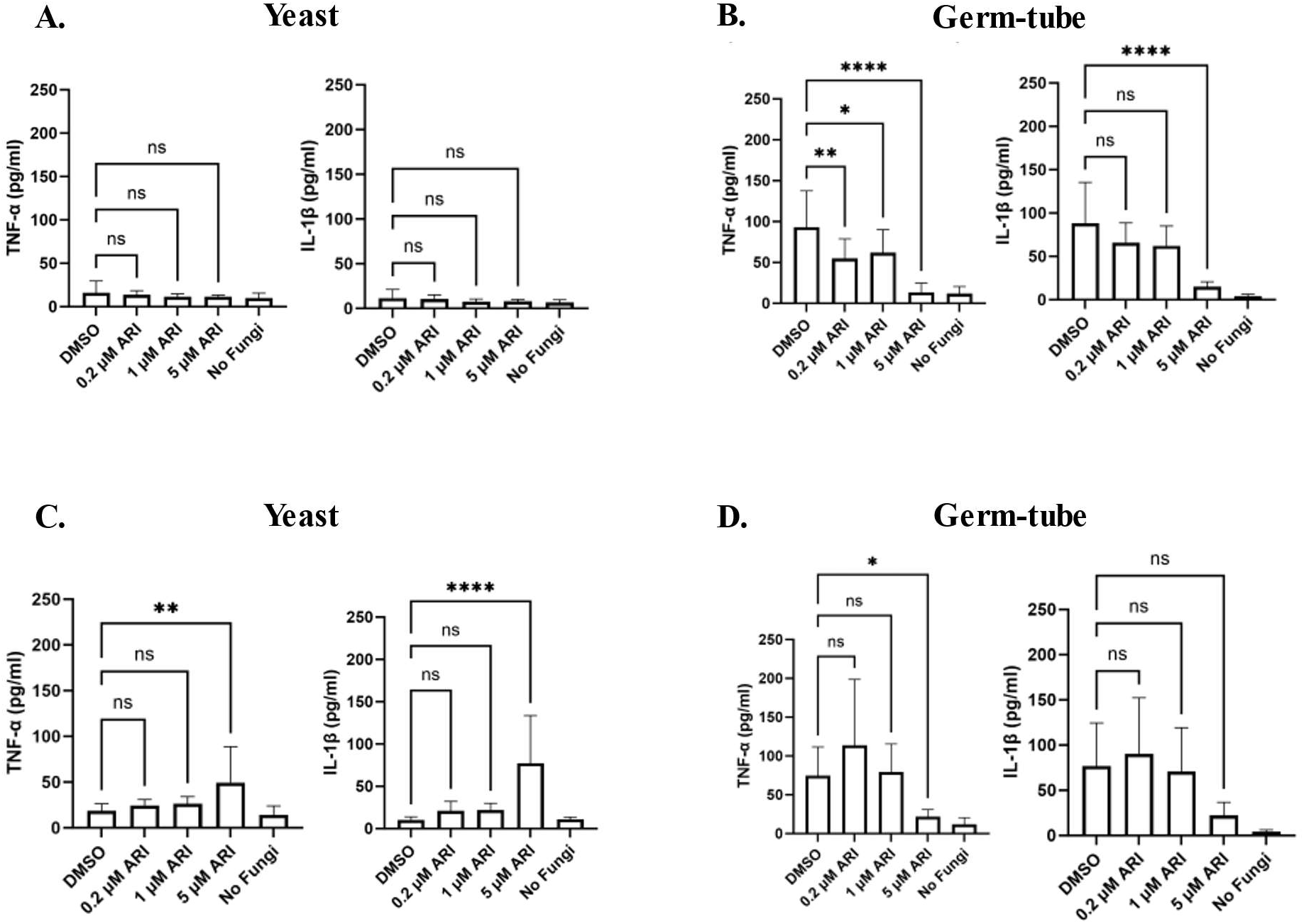
Aripiprazole alters *Candida albicans*-macrophage interaction. **A.** *C. albicans* strain SC5314 was grown in YNB-pH 7 medium supplemented with 0.2-5 µM aripiprazole (ARI) or 0.5% DMSO (vehicle control) for 24-hours at 3O°C to achieve yeast-form cells before being fixed in 10% formalin. Macrophages were co-incubated with fixed yeast at an MOI of 20:1 at 37°C for 24-hours and supernatant collected. ELISA assays were performed to quantify TNF-α or IL-lβ. Experiments were conducted in technical quadruplicate and performed independently in biological triplicate. Data is depicted as the mean + standard deviation. Statistical significance was calculated using one-way ANOVA test and Dunnet’s post-test. **B.** SC5314 was grown in RPMI-pH 7 medium supplemented with the same aripiprazole concentrations or vehicle control for 1.5-hours at 37°C to encourage germ-tube formation, then fixed as described in **(A.).** Macrophages were co-incubated with fixed germ-tube cells cells at an MOI of 20:1 at 37°C for 8-hours before supernatant collected. ELISA assays were performed as described in **(A.). C-D.** SC5314 was grown as yeast **(C.)** or germ-tubes **(D.)** and fixed as described above. Cells were added to macrophages at an MOI of 20:1 in the presence of 0.2-5 µM ARI or 0.5% DMSO. Macrophages were co-incubated as described in **(A.)** or **(B.).** TNF-α and IL-lβ release were quantified by ELISA. Data is depicted as the mean + standard deviation. Statistical significance was calculated using one-way ANOVA and Dunnet’s post-test. ***\* P <*** 0.01, ** ***P*** < 0.001, **** ***P <*** 0.0001. ns = not significant

## Discussion

Most infectious fungi are opportunists that principally cause disease in individuals with impaired immune function. Yet despite well-defined risk factors, the occurrence of IFIs in individual patients is difficult to predict, as are the clinical outcomes of subsequent therapeutic intervention. Paralleling this situation, acute and recurrent vulvovaginal candidiasis, have an idiopathic occurrence, with most affected women in otherwise good health (34–36). Each individual patient at risk of IFI presents a unique set of physiological, immunogenetic and clinical contexts including the regimen of medications consumed. Our long-term goal is to better understand if, and how, medications consumed by humans for indications unrelated to fungal infection (i.e. non-antifungal drugs) influence fungal physiology and pathogenicity, and if such drug-fungal interactions have the potential to affect disease initiation, progression, or resolution. Such drug-fungus interactions are pertinent, as patients at greatest risk of IFIs typically have one or more underlying disease condition that require treatment with complex regimens of drugs tailored to individual need. This includes those taken transiently and those intended for long-term or even life-long use, creating an additional complex and dynamic variable between at risk individuals. Furthermore, several species of *Candida* are natural residents of the gastrointestinal and reproductive tracts of healthy human subjects (37,38) and therefore are routinely exposed to medications administered to their human host. Certainly, drugs that cause profound immune dysfunction including corticosteroids and antineoplastic agents are known to dramatically increase a patient’s risk of developing an IFI (39). The use of broad-spectrum antibiotics that eliminate significant portions of the endogenous bacterial microbiota is also a significant risk-factor for oral and vaginal *Candida* infections (40). However, aside from those that have a direct and obvious effect upon the mammalian host’s primary defense mechanisms, the influence of most drugs upon the incidence and outcomes of invasive mycoses has received little attention. Several studies have sought to identify approved medications with stand-alone antifungal activity, or that potentiate the activity of existing antifungal agents, and can therefore potentially be repurposed to treat IFIs (12–14). Several studies also identified approved drugs that appear to antagonize or oppose the activity of antifungal therapeutics (15,41–43). However, the rudimentary outcomes of these studies (e.g. growth inhibition) are unlikely to identify those that have subtle or nuanced effects upon fungal physiology, pathogenicity, or immunogenicity that could potentially alter fungal fitness or interaction with the mammalian host and therefore the initiation, progression, or outcome of infection. Given the fundamental similarity of fungi and mammals at the molecular level, it is likely that many drugs designed to induce a biological response in human cells have an analogous impact upon fungal physiology. But the physiological consequences of exposing commensal and pathogenic fungi to most medications approved for human use remains unknown.

This specific study assessed the impact of several drugs we previously found to oppose the antifungal activity of the azole class of antifungals, upon *C. albicans* (16). Genetically encoded azole-resistance in *C. albicans* is often associated with point mutations in one or more key transcription factors that regulate the expression of the target enzyme Erg11p, or of multi-drug efflux pumps. However, mutations that confer antifungal resistance are often associated with fitness costs in the absence of the selecting agent (44,45), including point mutations that activate either Tac1p or Upc2p transcription factors (20,46). Therefore, it is surprising that despite six of the seven antagonists investigated in this study acting in a Tac1p-dependent manner, and the seventh in a Upc2p-dependent manner, only two had an obvious impact on *C. albicans* filamentous growth or phenotype at concentrations sufficient to substantially elevate azole MIC. Thus, azole-antagonistic activity of a drug or xenobiotic is not necessarily or inherently associated with significant detrimental impacts on *C. albicans* fitness. Consistent with our phenotypic data, the transcription of a surprisingly small number of genes was affected by all three of the antagonists examined. Further, many of the drug-responsive genes identified were consistent with a Tac1p-dependent mechanisms, including etofenamate, that we had previously shown to act largely through a Upc2p-dependent mechanism, indicating they affect a very specific aspect of fungal physiology. It is therefore likely that the majority of the 13 drugs we previously identified as fluconazole antagonists (16), also act through either Tac1p and/or Upc2p-dependent mechanisms, despite dissimilarity in structural, chemical, and physical properties. Viewed from this perspective, it appears *C. albicans* has evolved systems to specifically sense and respond to exogenous xenobiotics, which incidentally are adept at sensing medicinal compounds. One possibility is that as *C. albicans* has co-evolved with its mammalian host, these sensing mechanisms arose to detect host hormones or other signals that help the fungus regulate its physiology to favor commensalism or tissue invasion. A related prospect is that as part of a large and complex community of endogenous microbes residing within non-sterile body sites, xenobiotic detection systems may have also evolved to respond to competing microbes that release antifungal molecules. In these circumstances the seemingly promiscuous nature of the Tac1p and Upc2p-based xenobiotic response modules, as well as the broad-substrate specificities of the drug-efflux pumps regulated by Tac1p would be advantageous to respond to a multitude of distinct threats.

Much of this study focused upon the atypical antipsychotic aripiprazole to establish proof-of-principle - that previously unanticipated drug-fungus interactions have the potential to influence the outcome of invasive fungal disease. While its azole-antagonistic activity observed *in vitro* did not translate to reduced antifungal efficacy in the mouse model of disseminated infection, it expedites the progression of infection in the absence of antifungal therapy. Intuitively, this could be due to effect on the fungus that alter its inherent virulence, or upon the mammalian host that impact its intrinsic resistance to infection or its capacity to mount an effective immune response. Based on the observation that aripiprazole suppressed hyphal formation under *in vitro* conditions, and the importance of yeast-hypha morphogenesis to *C. albicans* virulence (47), worsened outcome with its administration the mouse model of infection is surprising. However, we did not observe any obvious effect of aripiprazole treatment upon *C. albicans* morphotype in tissue sections from infected mice. Thus, given the multitude of signaling pathways able to activate yeast-hypha morphogenesis (25), it is possible that aripiprazole’s effect upon hyphal growth is lessened by robust filamentation cues within mammalian tissue. An alternative possibility is that by delaying the transition of the yeast form provided in the initial inoculum into the more tissue invasive form, aripiprazole enhances the dispersal of infectious particles from initial foci of infection, leading to a greater number of active infection sites within the kidney and other tissues. Either way, the tissue lesions observed in *C. albicans* infected mice treated with aripiprazole are much larger, levels of kidney damage sustained greater, and the lesions are associated with elevated levels of immune cell infiltration compared to the mock treated controls.

To further examine this relationship, BALB/c mice were treated with aripiprazole as described previously, then infected with a yeast locked strain (CAI4+pKE4-NRG1). While this dose (4 x 10^5^ cells) did not induce lethality, aripiprazole treated infected mice displayed increased morbidity and kidney weights unrelated to fungal burden (fig. S5). Collectively, our data suggests worsened disease progression during aripiprazole treatment was independent of hyphal morphogenesis.

One possible explanation for this finding is that aripiprazole altered *C. albicans* immunogenicity in such a way as to promote a vigorous and potentially damaging host response. This could explain why the effects of aripiprazole were lost in the immunosuppressed models of disseminated candidiasis infection Notably, exposing *C. albicans* to sub-growth inhibitory concentrations of the echinocandin antifungals increases β-glucan exposure and immune recognition both *in vitro* and *in vivo* (48,49) (50,51). We were able to detect differences in levels of surface exposed chitin on aripiprazole treated *C. albicans*. However, chitin exposure is typically associated with fungal tolerance (50), and our *in vitro* studies indicate that fixed *C. albicans* cells pre-treated with aripiprazole stimulate less proinflammatory cytokine release from macrophages than untreated cells.

Although our studies were initiated following our observation that aripiprazole affects *C. albicans* physiology, we were not able to discount that the drug directly or indirectly affected the hosts response to infection, resulting less effective defense. Curiously, aripiprazole also appears to affect the human macrophage responses, dampening release of two key pro-inflammatory cytokines in response to fixed *C. albicans*. Thus, it is possible aripiprazole may exert both drug-fungus and drug-host interactions that impact the outcome of invasive *C. albicans*. Indeed, aripiprazole use in humans has also been associated with decreased pro-inflammatory cytokine levels (52). Based on previous studies, both TNF-α and IL1-β are expected to have a protective role in disseminated *C. albicans* infection (53–55), thus it is not surprising that suppression of its release from key cells of the innate immune response would have a detrimental impact on the outcome of infection in mice. Nonetheless, at first glance, the suppression of key pro-inflammatory responses by macrophages is inconsistent with the apparently hyperinflammatory lesions observed in aripiprazole treated mice infected with *C. albicans*. These findings could be reconciled by the inherent complexity and multi-faceted nature of the mechanisms by which mammals detect and respond to infectious fungi. Hematopoietic, epithelial, and endothelial cells all make important contributions to host defense against pathogenic fungi (50), with each responding to a discrete cohort of PAMPs and often producing qualitatively or quantitatively distinct responses. For example, macrophages preferentially respond to *C. albicans* hyphal over yeast forms to produce pro-inflammatory cytokines (56), peripheral blood mononuclear cells (PBMCs) on the other hand respond more vigorously to the yeast form (57). Thus, it is possible the very same aripiprazole induced changes in PAMP presentation that suppress macrophage responses, could hyper-stimulate pro-inflammatory responses in other host cell types. Either way, both our *in vitro* and *in vivo* data clearly establish that aripiprazole changes the nature of the *C. albicans*-host interaction in ways that can affect disease progression.

In a broader context, this study raises the issue of biologically active molecules including the medications consumed by colonized or infected individuals disrupting either mammalian or fungal physiology, and thus potentially the equilibrium of the host-pathogen interaction. Drug-fungus interactions for example could alter the outcome of infection by: 1). Causing profound physiological dysfunction that compromises the viability, fitness, or pathogenicity of the fungus *in vivo*; 2). Changing the surface characteristics of the fungus in ways that alter its immunogenicity, binding/activation of complement, or sensitivity to host derived antimicrobial peptides; 3). Altering fungal sensitivity to antifungal drugs. It is also important to consider the effects of each drug at therapeutically relevant drug concentrations, which in the context of disseminated infection, most investigators will take as unbound blood serum concentrations taken from patients on a standard dosing regimen. However, *Candida* species naturally colonize the GI tract, where the concentration of orally administered drugs (or those excreted via the GI tract) is potentially much higher. Additionally, drug concentrations within specific tissues can differ markedly from serum, and extensive tissue dissolution within foci of infection could have further affects. We observed *C. albicans* infected mice had significantly higher serum concentrations of aripiprazole (∼1500 ng/ml) than uninfected mice treated with the same dosing (∼450 ng/ml) (fig. S6). This is most likely because aripiprazole is almost entirely removed by metabolism in the liver, and often genes encoding drug metabolizing enzymes are downregulated in conditions in systemic inflammation and similar stress (like acute infection) (58–61). This could lead to less drug metabolism, leading to increased plasma concentration. Such affects can further modulate host response as well as fungal physiology.

The effect of any individual medication on the outcome of IFIs is unlikely to be immediately apparent in the clinical setting due to the intricacies of each infected individuals’ circumstances that include complex and highly varied drug regimens provided and relatively small numbers of infected patients treated with any given medication. Additionally, the effects of drug-fungus or even drug-host interactions upon the course of an IFI is also likely to be context dependent, with many factors such as the causative species, specific nature and severity of a patient’s immune deficiency, site of infection, and potentially genetic factors being key determinants. Thus, it remains to be determined how many drugs approved for human use have the potential to modulate the outcome of IFIs. Nonetheless, the findings of this study can serve to raise awareness of the potential influence of non-antifungal medications on the outcome of invasive fungal infections.

## Acknowledgements.

Research reported in this publication was supported by the National Institute of Allergy and Infectious Diseases of the National Institutes of Health under Award Number R01AI152067 and R33AI127607 (to GEP) and R01AI134796 (to BMP). The content is solely the responsibility of the authors and does not necessarily represent the official views of the National Institutes of Health.

---

**Figure S1.**
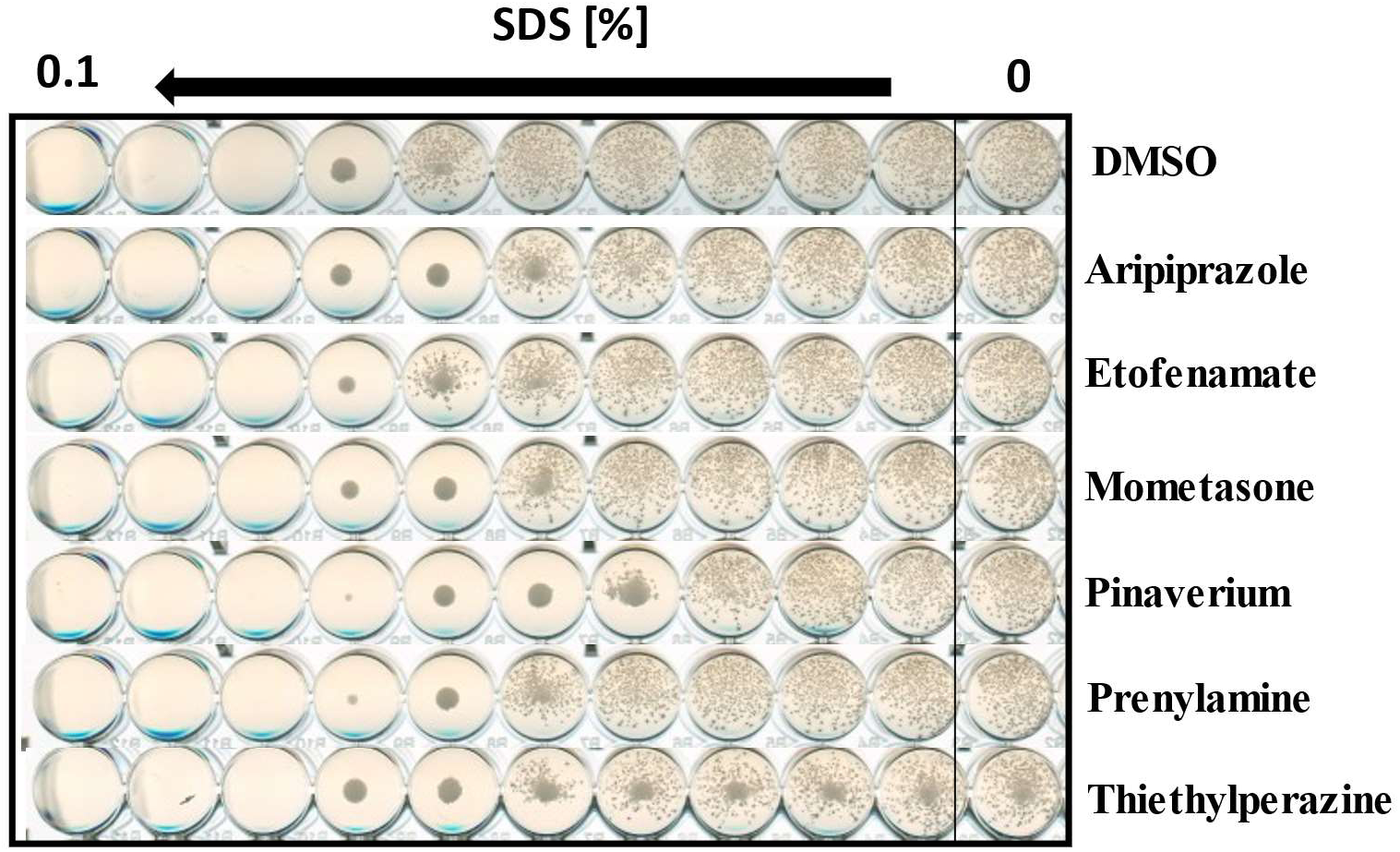
Several antifungal antagonists decrease tolerance to the cell membrane stressor SDS. Stressor susceptibility assays were set conducted with *C. albicans* SC5314 in 96-well plates at 1 x 10^4^ cells/ml in RPMI-pH 7 containing increasing 2-fold concentrations of the cell membrane stressor SDS. Cells were grown in the presence of either DMSO or 5 µM of each medication, and then incubated at 35°C. Plates were imaged at 24-hours.

**Figure S2.**
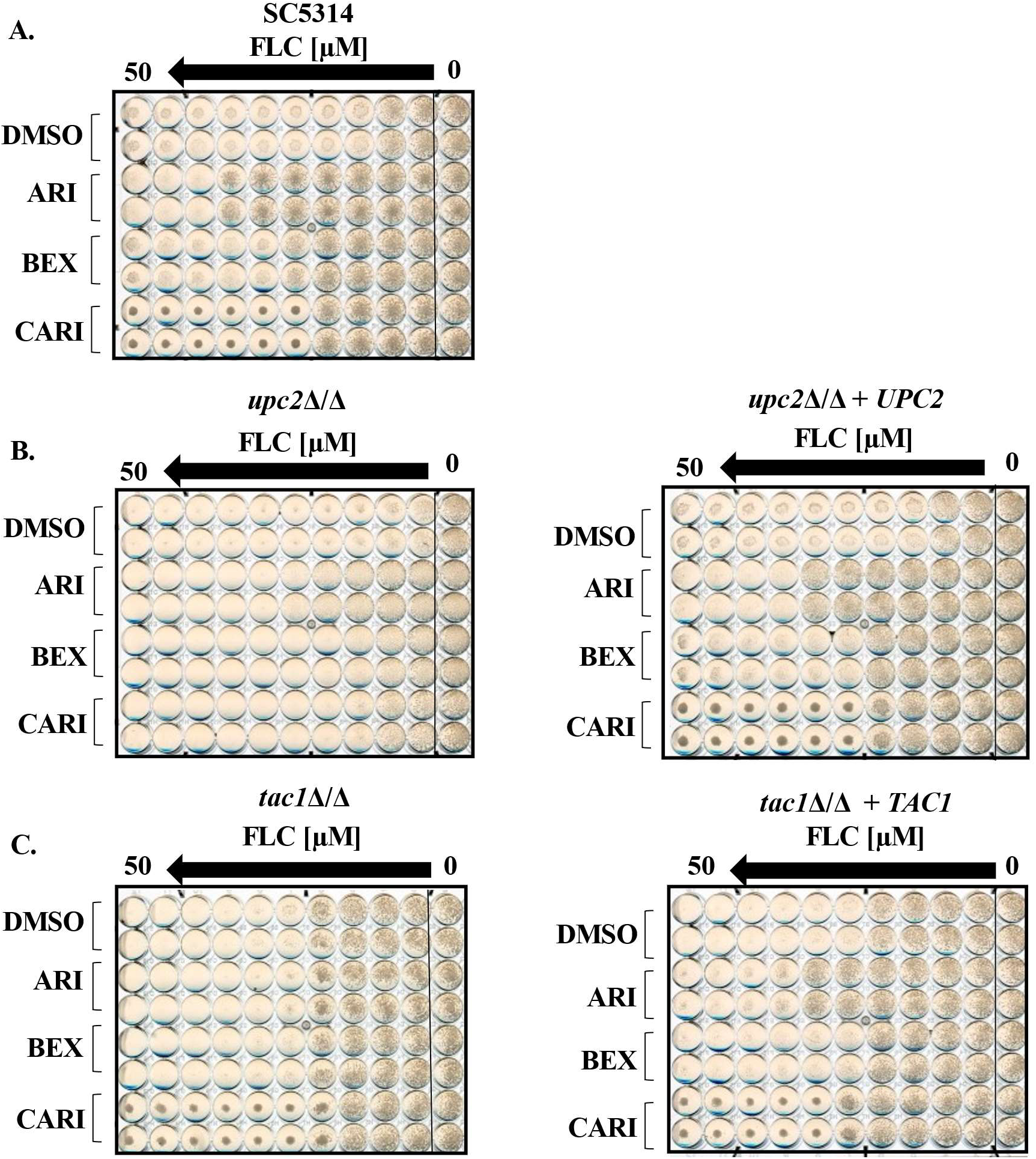
Aripiprazole analogs act through distinct mechanisms. **A.** Fluconazole (FLC) susceptibility assays were performed with *C.albicans* strain SC5314 using the CLSI method. Medium was supplemented with either 5 μM aripiprazole (ARI), bexpiprazole (BEX), cariprazine (CARI), or 0.5% DMSO (vehicle control) and imaged after 24-hours of incubation. Plates are representative of assays performed in biological duplicate. **B-C.** Fluconazole susceptibility assays were performed as describd above with a *upc2*Δ/Δ strain and *upc2*Δ/Δ + *UPC2* derived strain (**B.**) or *tac1*Δ/Δ and its *tac1*Δ/Δ + *TAC1* derived strain accordingly (**C.**). Plates are representative of assays performed in biological duplicate.

**Figure S3.**
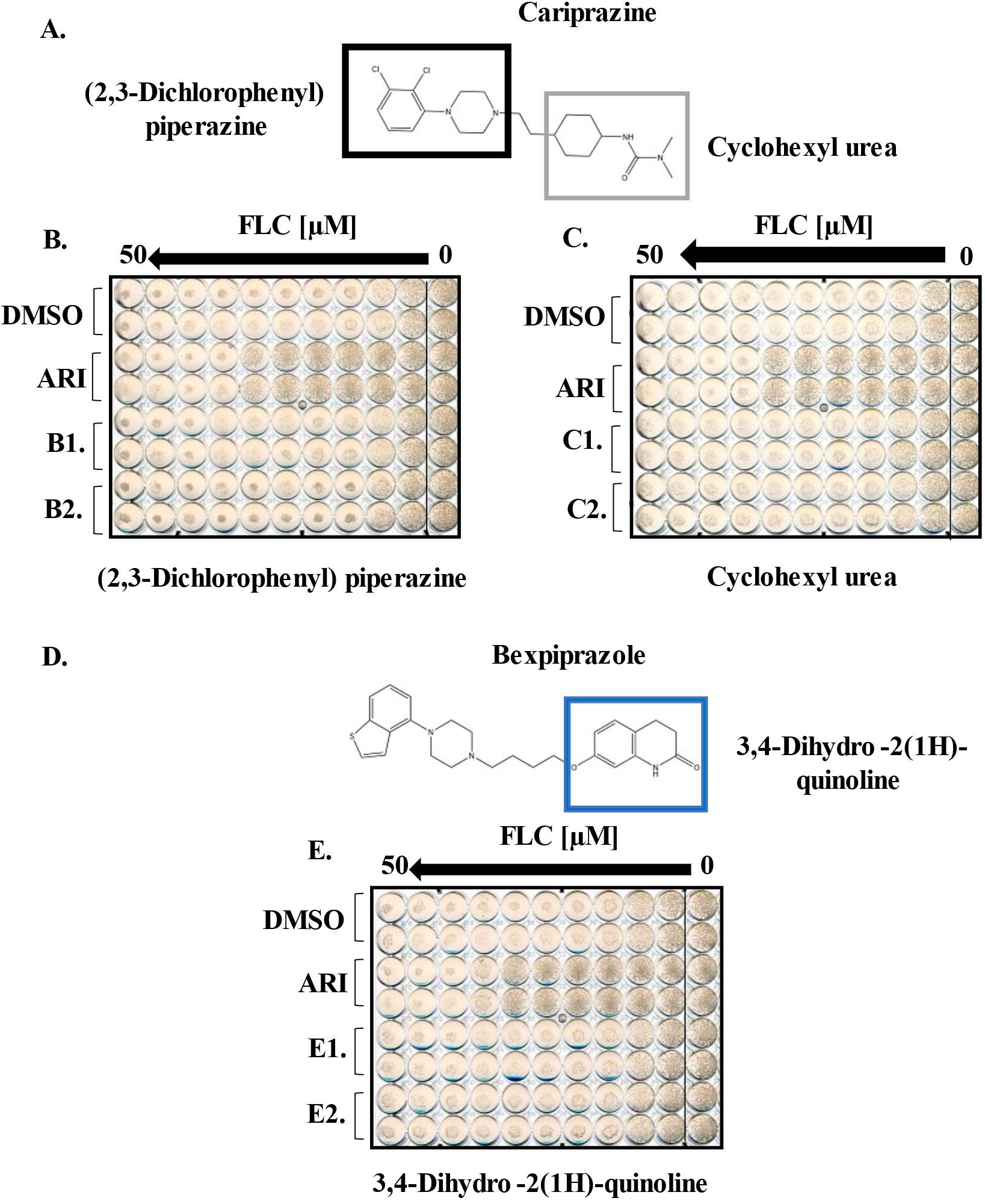
Cariprazine and bexpiprazole sub-structures alone are insufficient to alter fluconazole activity against *Candida albicans.* **A.** The chemical structure of cariprazine with the two sub-structures highlighted: (2,3-diclτlorophenyl)piperazine (indicated by a black square) or cyclohexyl urea (indicated by a gray square). **B.** Fluconazole (FLC) susceptibility assays were performed with *C. albicans* strain SC5314 using CLSI conduct. Medium was supplemented with either 0.5% (DMSO) 5 µM aripiprazole (ARI), 5 µM **(BI.),** or 25 µM **(B2.)** of (2,3-Dichlorophenyl) piperazine and plates imaged after 24-hours of incubation. Plates are representative of assays performed in biological duplicate. **C.** FLC susceptibility assays were performed as described in **(B.),** using the substructure cyclohexyl at two concentrations: 5 µM **(Cl.)** or 25 µM **(C2). D.** Bexpiprazole containing the sub-structure 3,4-Dihydro-2(lH)-quinoline (indicated by a blue square). **E.** FLC susceptibility assays were performed as described in **(B.),** with using the substructure 3.4-dihydro-2(lH)-quinoline at 5 µM **(El.)** or 25 µM **(E2.).**

**Figure S4.**
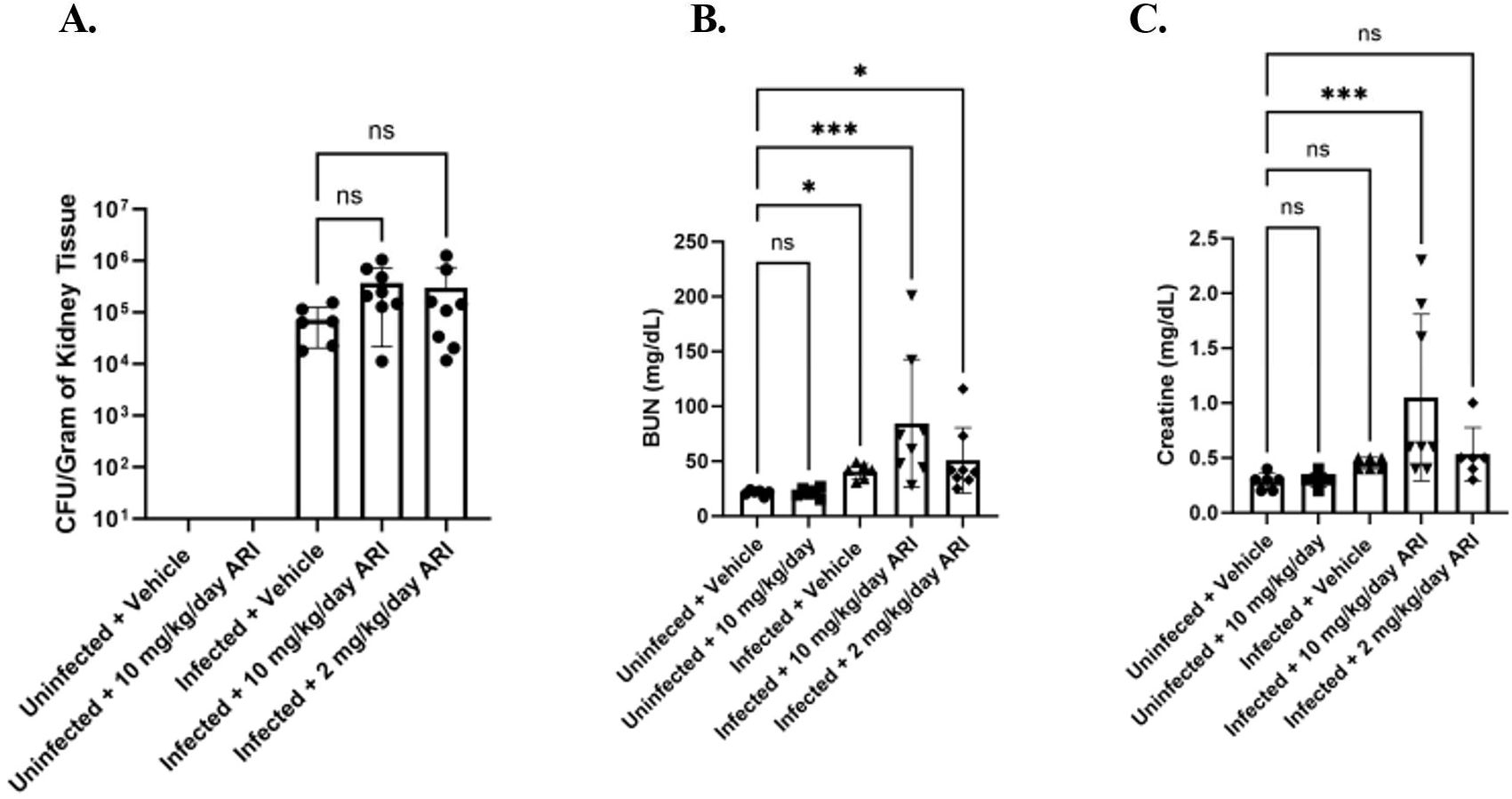
Aripiprazole effects disease progression in C57BL/6 mice. **A.** Groups of C57BL/6 mice (n = 8) were treated daily with either 10 or 2 mg/kg/day aripiprazole or vehicle starting 3 days prior to infection and then infected with 2.5 x 10^5^ of SC5314 or PBS (vehicle control). At 48-hour endpoints, kidneys were excised, and fungal burden quantified as CFU per gram of tissue. Data is depicted as the mean + standard deviation. Statistical significance was calculated using a one-way ANOVA and Kruskal-Wallis post-test. **B-C.** Blood was extracted from the same mice as in **(A.)** serum processed, and blood urea nitrogen (BUN) **(B.)** or creatinine levels **(C.)** were quantified. Groups were compared to uninfected vehicle group. Data is depicted as the mean + standard deviation, and statistical significance was determined using a one-way ANOVA and Kruskal-Wallis post-test (n = 8).

**Figure S5:**
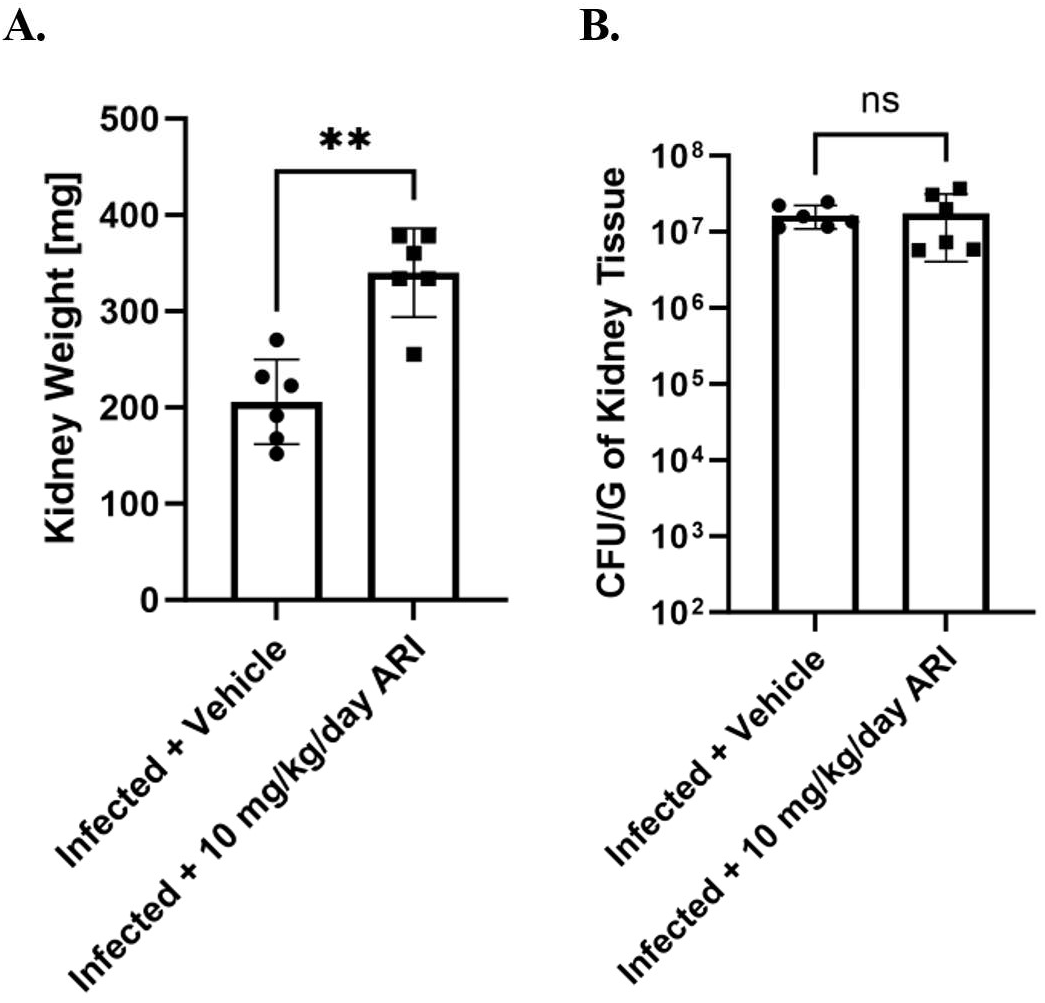
Aripiprazole exacerbates disease in a disseminated candidiasis infection. **A.** Groups of BALB/c mice (n = 6) were treated daily with 10 mg/kg/day aripiprazole (ARI) or vehicle starting 3 days prior to infection. Mice were then infected with 4 x 10^5^ of the CAI4+pKE4-NRG 1 yeast-locked strain, and 5 days p.i, mice were euthanized, kidneys were excised. Kidney weight was recorded, and weight was compared between drug and vehicle heated groups. Data is depicted as the mean + standard deviation, and statistical significance calculated using a one-way ANOVA. and Mann-Whitney post-test. B. Kidneys from mice in **(A.)** were homogenized, and fungal burden was quantified as CFU per gram of tissue. Data is depicted as the mean + standard deviation, and statistical significance calculated using a one-way ANOVA and Mann-Whitney post-test. *\*\* P<* 0.005. ns = not significant

**Figure S6:**
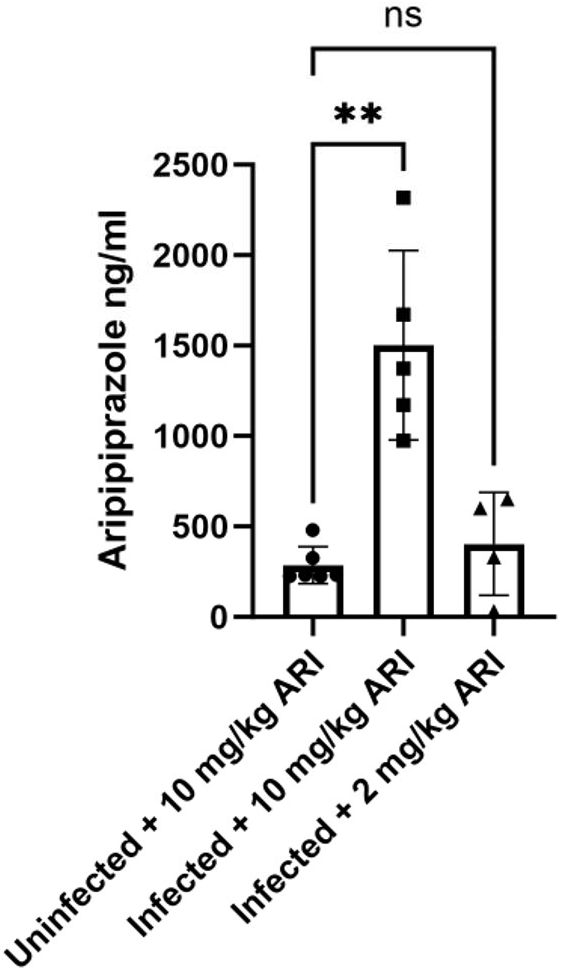
Aripiprazole concentration increases during disseminated candidiasis infection. Groups of BALB/c mice (n = 6) were treated daily with 10 or 2 mg/kg/day aripiprazole (ARI) starting 3 days prior to infection. Mice were then infected with 2.5 x 10^5^ of SC5314, and drug treatment continued. At the 48-hour endpoint, blood was collected, plasma extracted, and aripiprazole concentration was quantified using LC-MS. Infected drug treated mice were compared to uninfected treated mice. Data is depicted as the mean + standard deviation, and statistical significance was calculated using a one-way ANOVA and Kruskal-Wallis post-test. ***\*\* P <*** 0.005. ns = not significant

**Table S1.**
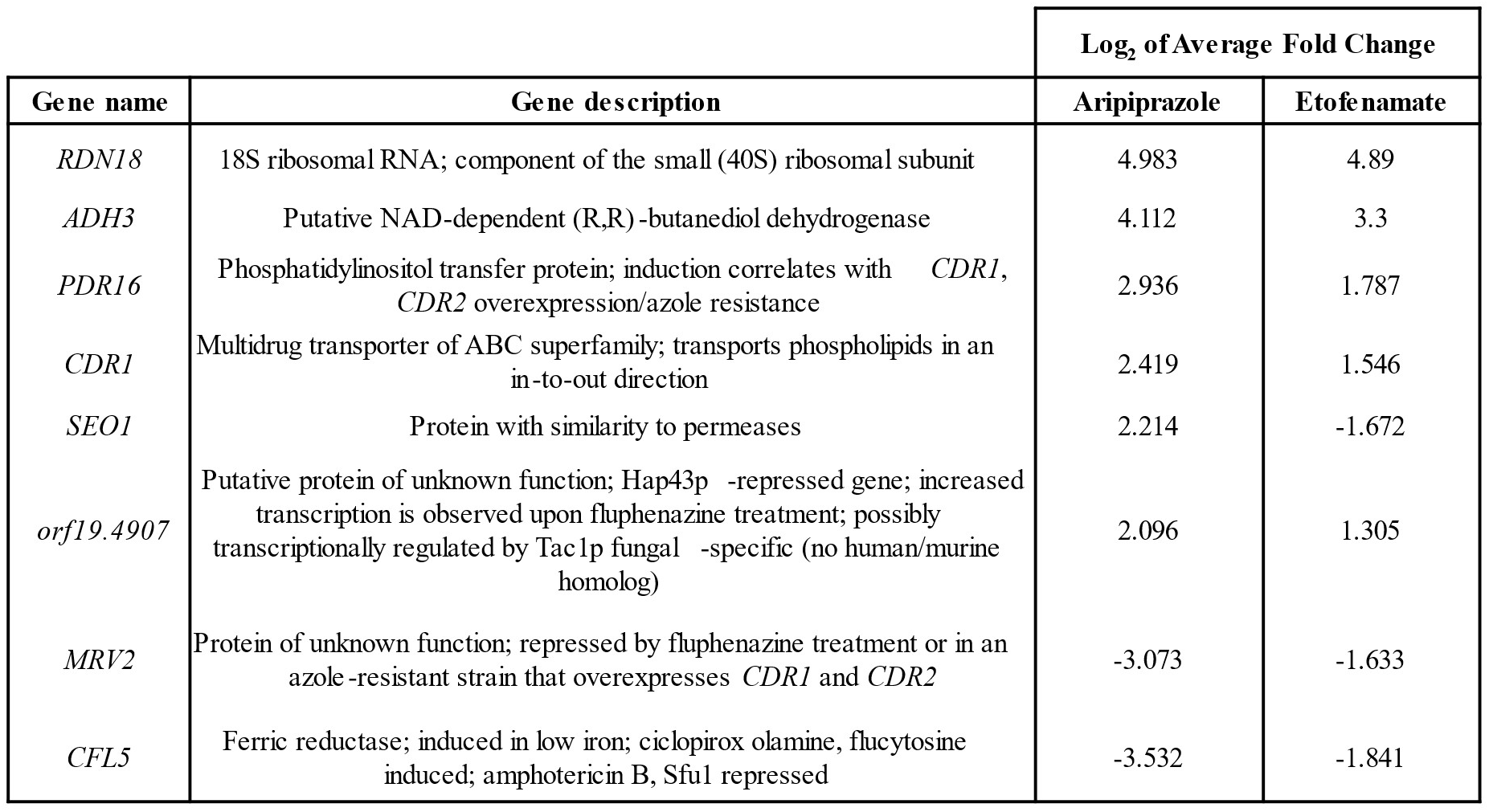
Overlap of aripiprazole and etofenamate responsive genes. *C. albicans* was grown in RPMI-pH 7 at 35^θ^C for 6-hours with 5 µM of aripiprazole, etofenamate, or 0.5% DMSO (vehicle). High throughput sequence analysis was performed, and significantly altered (log_2_fold > 1 or < -1, adjusted to p < 0.5) gene expression (compared to vehicle controls) determined.

| Gene name | Gene description | Log<sub>2</sub> of Average Fold Change |  |
| --- | --- | --- | --- |
|  |  | Aripiprazole | Etofenamate |
| *RDN18* | 18S ribosomal RNA; component of the small (40S) ribosomal subunit | 4.983 | 4.89 |
| *ADH3* | Putative NAD-dependent (R,R)-butanediol dehydrogenase | 4.112 | 3.3 |
| *PDR16* | Phosphatidylinositol transfer protein; induction correlates with *CDR1*, *CDR2* overexpression/azole resistance | 2.936 | 1.787 |
| *CDR1* | Multidrug transporter of ABC superfamily; transports phospholipids in an in-to-out direction | 2.419 | 1.546 |
| *SEO1* | Protein with similarity to permeases | 2.214 | -1.672 |
| *orf19.4907* | Putative protein of unknown function; Hap43p -repressed gene; increased transcription is observed upon fluphenazine treatment; possibly transcriptionally regulated by Tac1p fungal -specific (no human/murine homolog) | 2.096 | 1.305 |
| *MRV2* | Protein of unknown function; repressed by fluphenazine treatment or in an azole-resistant strain that overexpresses *CDR1* and *CDR2* | -3.073 | -1.633 |
| *CFL5* | Ferric reductase; induced in low iron; ciclopirox olamine, flucytosine induced; amphotericin B, Sfu1 repressed | -3.532 | -1.841 |

**Table S2.**
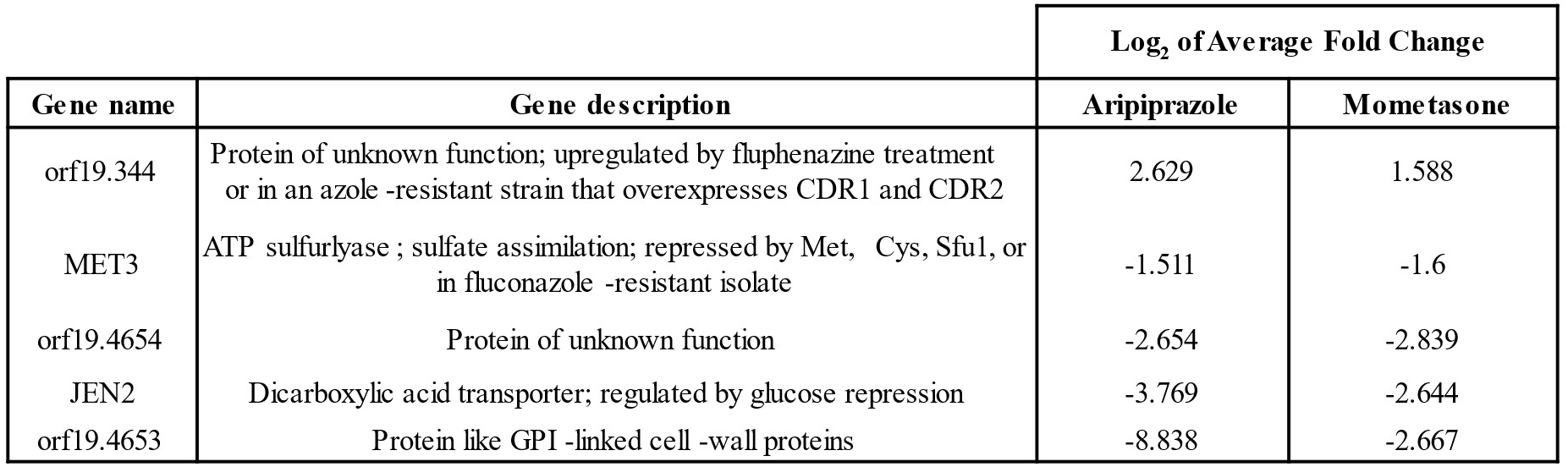
Overlap of aripiprazole and mometasone responsive genes. *C. albicans* was grown in RPMI-pH 7 at 35°C for 6-hours with 5 µM of aripiprazole, mometasone, or 0.5% DMSO (vehicle). High throughput sequence analysis was performed, and significantly altered (log_2_fold > 1 or < -1, adjusted to p < 0.5) gene expression (compared to vehicle controls) determined.

**Table S3.** Overlap of etofenamate and mometasone responsive genes. *C. albicans* was grown in RPMI-pH 7 at 35°C for 6-hours with 5 µM of etofenamate, mometasone, or 0.5% DMSO (vehicle). High throughput sequence analysis was performed, and significantly altered (log_2_fold > 1 or < -1, adjusted to p < 0.5) gene expression (compared to vehicle controls) determined.

| Gene name | Gene description | Log<sub>2</sub> of Average Fold Change |  |
| --- | --- | --- | --- |
|  |  | Etofenamate | Mometasone |
| *snR57b* | C/D box small nucleolar RNA (snoRNA) | -1.244 | -1.606 |

**Table S4.**
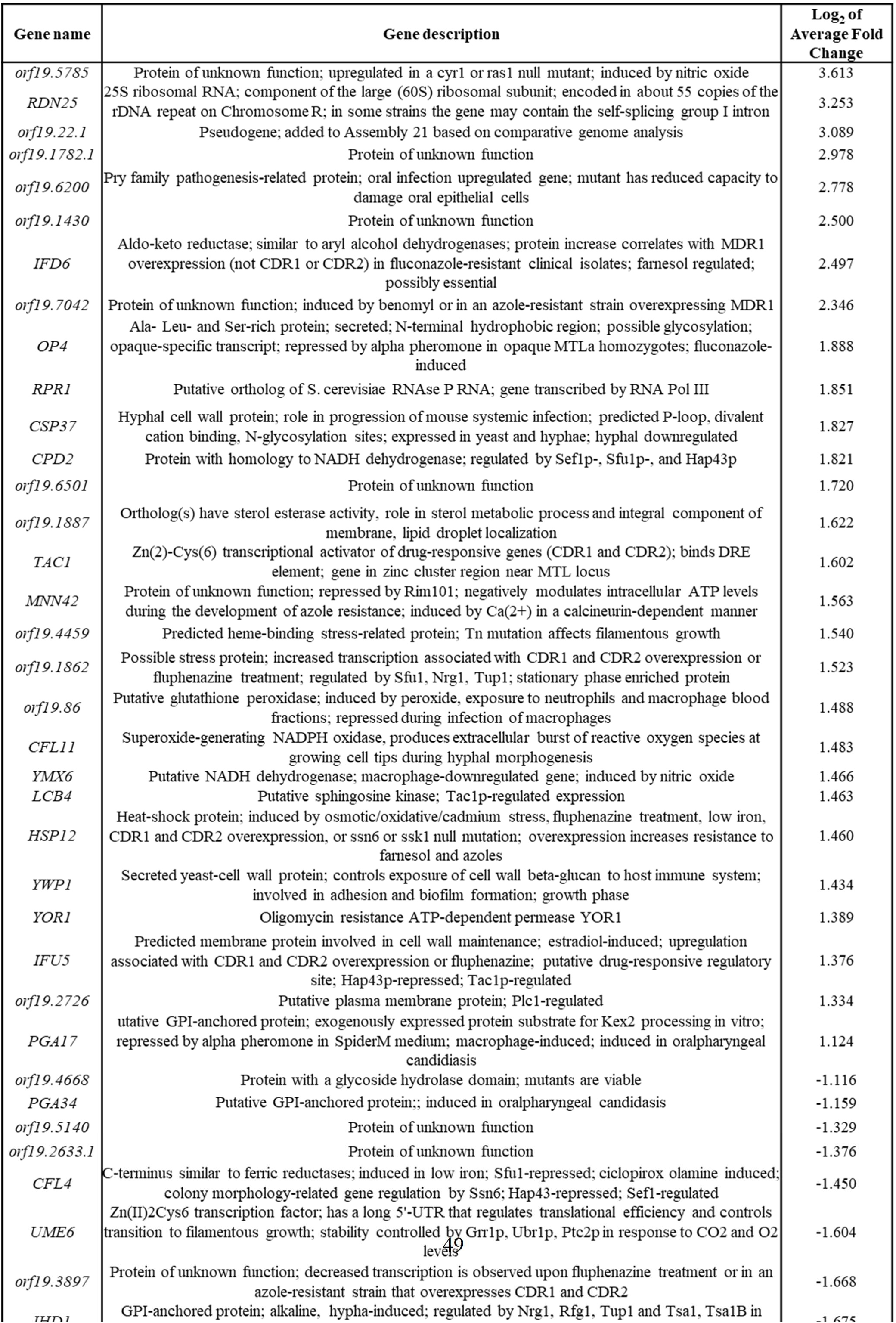
Genes responsive to aripiprazole only. *C. albicans* was grown in RPMI-pH 7 at 35°C for 6-hours with 5 µM of aripiprazole or 0.5% DMSO (vehicle). High throughput sequence analysis was performed, and significantly altered (log_2_fold > 1 or < -1, adjusted to p < 0.5) gene expression (compared to vehicle controls) determined.

| Gene name | Gene description | Log<sub>2</sub> of Average Fold Change |
| --- | --- | --- |
| *orf19.5785* | Protein of unknown function; upregulated in a *cyr1* or *ras1* null mutant; induced by nitric oxide | 3.613 |
| *RDN25* | 25S ribosomal RNA; component of the large (60S) ribosomal subunit; encoded in about 55 copies of the rDNA repeat on Chromosome R; in some strains the gene may contain the self-splicing group I intron | 3.253 |
| *orf19.22.1* | Pseudogene; added to Assembly 21 based on comparative genome analysis | 3.089 |
| *orf19.1782.1* | Protein of unknown function | 2.978 |
| *orf19.6200* | Pry family pathogenesis-related protein; oral infection upregulated gene; mutant has reduced capacity to damage oral epithelial cells | 2.778 |
| *orf19.1430* | Protein of unknown function | 2.500 |
| *IFD6* | Aldo-keto reductase; similar to aryl alcohol dehydrogenases; protein increase correlates with MDR1 overexpression (not CDR1 or CDR2) in fluconazole-resistant clinical isolates; farnesol regulated; possibly essential | 2.497 |
| *orf19.7042* | Protein of unknown function; induced by benomyl or in an azole-resistant strain overexpressing MDR1 | 2.346 |
| *OP4* | Ala- Leu- and Ser-rich protein; secreted; N-terminal hydrophobic region; possible glycosylation; opaque-specific transcript; repressed by alpha pheromone in opaque MTLA homozygotes; fluconazole-induced | 1.888 |
| *RPR1* | Putative ortholog of *S. cerevisiae* RNase P RNA; gene transcribed by RNA Pol III | 1.851 |
| *CSP37* | Hyphal cell wall protein; role in progression of mouse systemic infection; predicted P-loop, divalent cation binding, N-glycosylation sites; expressed in yeast and hyphae; hyphal downregulated | 1.827 |
| *CPD2* | Protein with homology to NADH dehydrogenase; regulated by Sef1p-, Sfu1p-, and Hap43p | 1.821 |
| *orf19.6501* | Protein of unknown function | 1.720 |
| *orf19.1887* | Ortholog(s) have sterol esterase activity, role in sterol metabolic process and integral component of membrane, lipid droplet localization | 1.622 |
| *TAC1* | Zn(2)-Cys(6) transcriptional activator of drug-responsive genes (CDR1 and CDR2); binds DRE element; gene in zinc cluster region near MTL locus | 1.602 |
| *MNN42* | Protein of unknown function; repressed by Rim101; negatively modulates intracellular ATP levels during the development of azole resistance; induced by Ca(2+) in a calcineurin-dependent manner | 1.563 |
| *orf19.4459* | Predicted heme-binding stress-related protein; Tn mutation affects filamentous growth | 1.540 |
| *orf19.1862* | Possible stress protein; increased transcription associated with CDR1 and CDR2 overexpression or fluphenazine treatment; regulated by Sfu1, Nrg1, Tup1; stationary phase enriched protein | 1.523 |
| *orf19.86* | Putative glutathione peroxidase; induced by peroxide, exposure to neutrophils and macrophage blood fractions; repressed during infection of macrophages | 1.488 |
| *CFL11* | Superoxide-generating NADPH oxidase, produces extracellular burst of reactive oxygen species at growing cell tips during hyphal morphogenesis | 1.483 |
| *YMX6* | Putative NADH dehydrogenase; macrophage-downregulated gene; induced by nitric oxide | 1.466 |
| *LCB4* | Putative sphingosine kinase; Tac1p-regulated expression | 1.463 |
| *HSP12* | Heat-shock protein; induced by osmotic/oxidative/cadmium stress, fluphenazine treatment, low iron, CDR1 and CDR2 overexpression, or *ssn6* or *ssk1* null mutation; overexpression increases resistance to farnesol and azoles | 1.460 |
| *YWP1* | Secreted yeast-cell wall protein; controls exposure of cell wall beta-glucan to host immune system; involved in adhesion and biofilm formation; growth phase | 1.434 |
| *YOR1* | Oligomycin resistance ATP-dependent permease YOR1 | 1.389 |
| *IFU5* | Predicted membrane protein involved in cell wall maintenance; estradiol-induced; upregulation associated with CDR1 and CDR2 overexpression or fluphenazine; putative drug-responsive regulatory site; Hap43p-repressed; Tac1p-regulated | 1.376 |
| *orf19.2726* | Putative plasma membrane protein; Plc1-regulated | 1.334 |
| *PGA17* | Putative GPI-anchored protein; exogenously expressed protein substrate for Kex2 processing in vitro; repressed by alpha pheromone in SpiderM medium; macrophage-induced; induced in oralpharyngeal candidiasis | 1.124 |
| *orf19.4668* | Protein with a glycoside hydrolase domain; mutants are viable | -1.116 |
| *PGA34* | Putative GPI-anchored protein; induced in oralpharyngeal candidiasis | -1.159 |
| *orf19.5140* | Protein of unknown function | -1.329 |
| *orf19.2633.1* | Protein of unknown function | -1.376 |
| *CFL4* | C-terminus similar to ferric reductases; induced in low iron; Sfu1-repressed; ciclopirox olamine induced; colony morphology-related gene regulation by Ssn6; Hap43-repressed; Sef1-regulated | -1.450 |
| *UME6* | Zn(II)2Cys6 transcription factor; has a long 5'-UTR that regulates translational efficiency and controls transition to filamentous growth; stability controlled by Grr1p, Ubr1p, Ptc2p in response to CO2 and O2 levels | -1.604 |
| *orf19.3897* | Protein of unknown function; decreased transcription is observed upon fluphenazine treatment or in an azole-resistant strain that overexpresses CDR1 and CDR2 | -1.668 |
| *IFD1* | GPI-anchored protein; alkaline, hypha-induced; regulated by Nrg1, Rfg1, Tup1 and Tsa1, Tsa1B in | -1.675 |

**Table S5.**
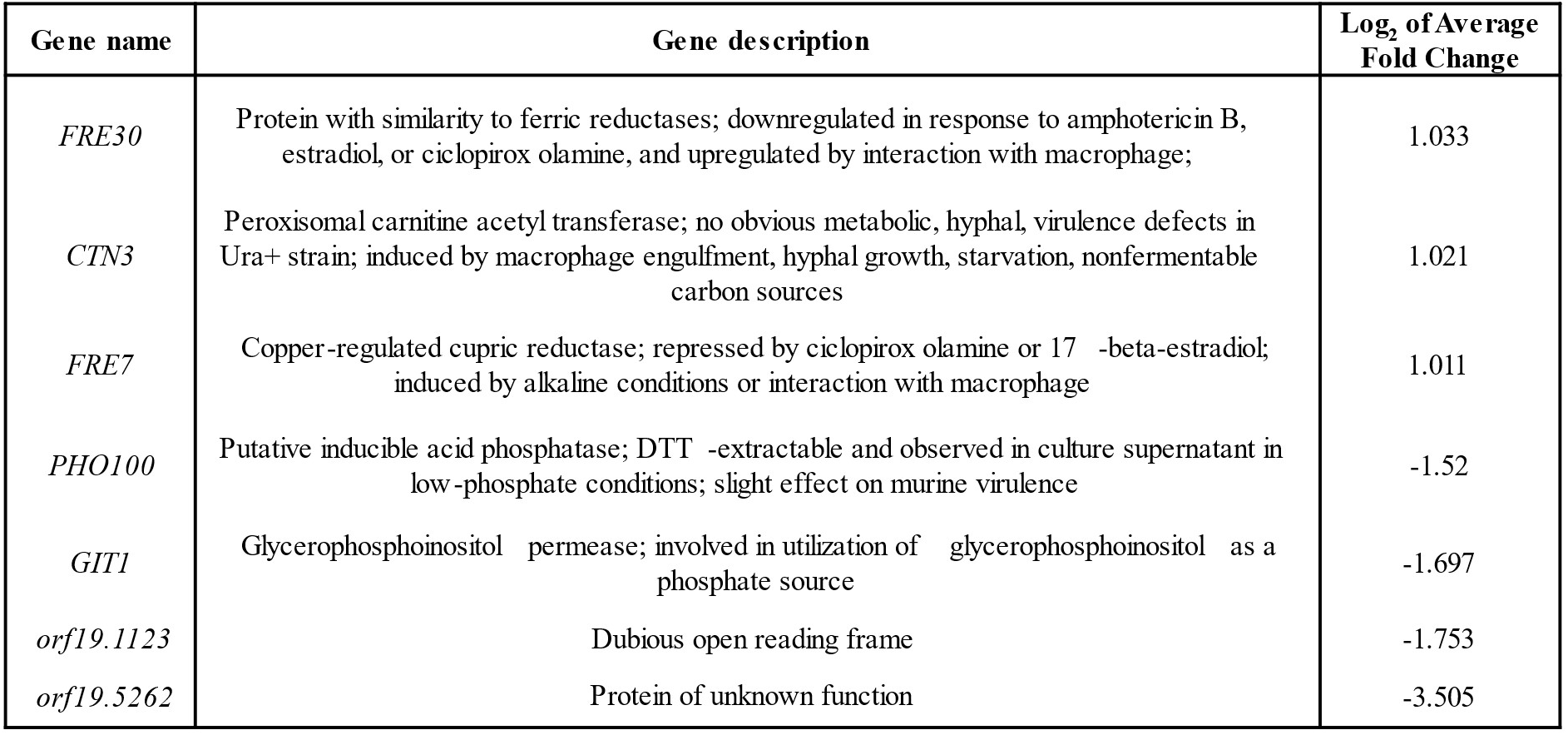
Genes responsive to etofenamate only. *C. albicans* was grown in RPMI-pH 7 at 35°C for 6-hours with 5 µM of etofenamate or 0.5% DMSO (vehicle). High throughput sequence analysis was performed, and significantly altered (log_2_fold > 1 or < -1, adjusted to p < 0.5) gene expression (compared to vehicle controls) determined.

| Gene name | Gene description | Log<sub>2</sub> of Average Fold Change |
| --- | --- | --- |
| *FRE30* | Protein with similarity to ferric reductases; downregulated in response to amphotericin B, estradiol, or ciclopirox olamine, and upregulated by interaction with macrophage; | 1.033 |
| *CTN3* | Peroxisomal carnitine acetyl transferase; no obvious metabolic, hyphal, virulence defects in Ura+ strain; induced by macrophage engulfment, hyphal growth, starvation, nonfermentable carbon sources | 1.021 |
| *FRE7* | Copper-regulated cupric reductase; repressed by ciclopirox olamine or 17 β-estradiol; induced by alkaline conditions or interaction with macrophage | 1.011 |
| *PHO100* | Putative inducible acid phosphatase; DTT -extractable and observed in culture supernatant in low -phosphate conditions; slight effect on murine virulence | -1.52 |
| *GIT1* | Glycerophosphoinositol permease; involved in utilization of glycerophosphoinositol as a phosphate source | -1.697 |
| *orf19.1123* | Dubious open reading frame | -1.753 |
| *orf19.5262* | Protein of unknown function | -3.505 |

**Table S6.**
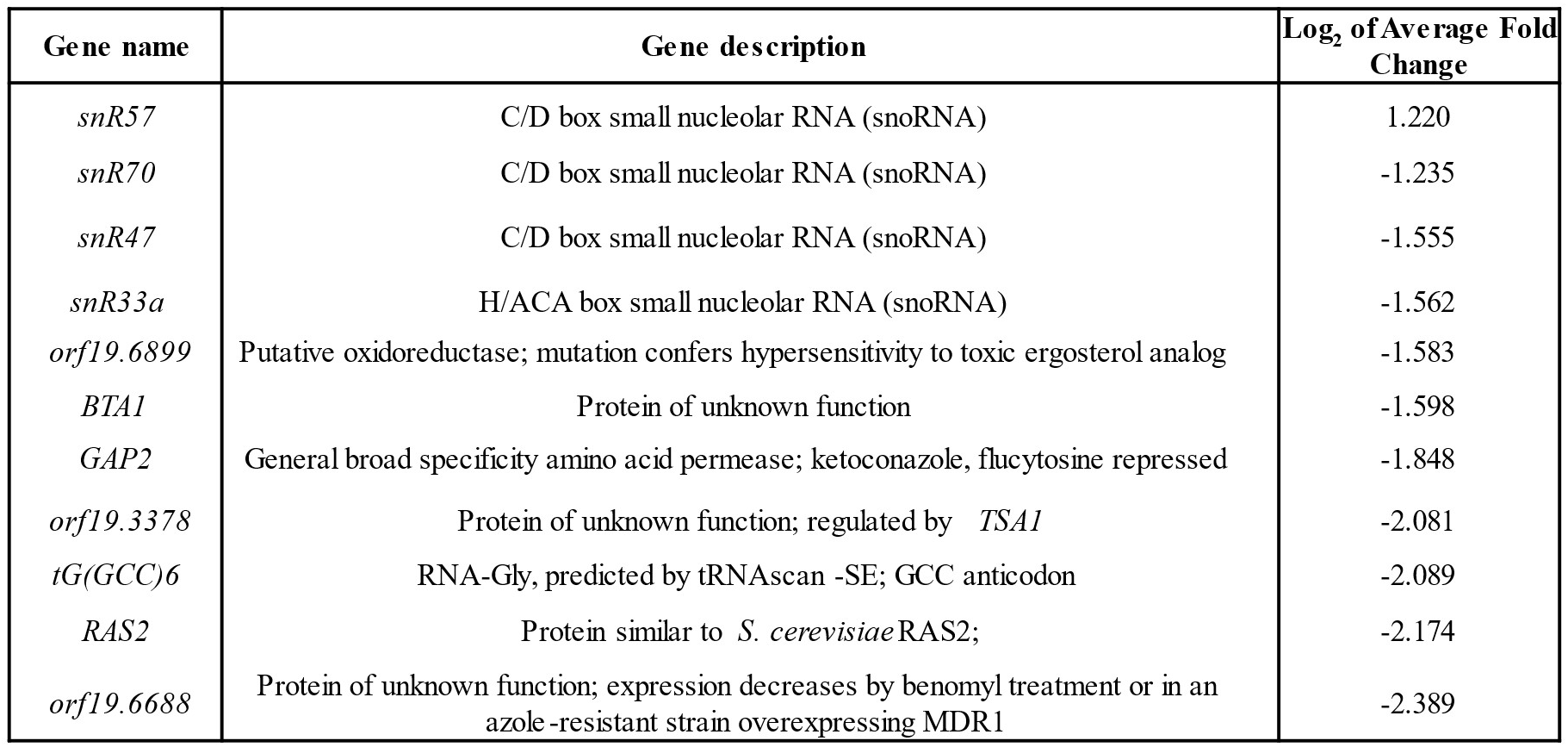
Genes responsive to mometasone only. *C. albicans* was grown in RPMI-pH 7 at 35^θ^C for 6-hours with 5 µM of mometasone or 0.5% DMSO (vehicle). High throughput sequence analysis was performed, and significantly altered (log_2_fold > 1 or < -1, adjusted to p < 0.5) gene expression (compared to vehicle controls) determined.

| Gene name | Gene description | Log<sub>2</sub> of Average Fold Change |
| --- | --- | --- |
| *snR57* | C/D box small nucleolar RNA (snoRNA) | 1.220 |
| *snR70* | C/D box small nucleolar RNA (snoRNA) | -1.235 |
| *snR47* | C/D box small nucleolar RNA (snoRNA) | -1.555 |
| *snR33a* | H/ACA box small nucleolar RNA (snoRNA) | -1.562 |
| *orf19.6899* | Putative oxidoreductase; mutation confers hypersensitivity to toxic ergosterol analog | -1.583 |
| *BTA1* | Protein of unknown function | -1.598 |
| *GAP2* | General broad specificity amino acid permease; ketoconazole, flucytosine repressed | -1.848 |
| *orf19.3378* | Protein of unknown function; regulated by *TSA1* | -2.081 |
| *tG(GCC)6* | RNA-Gly, predicted by tRNAscan -SE; GCC anticodon | -2.089 |
| *RAS2* | Protein similar to *S. cerevisiae* RAS2; | -2.174 |
| *orf19.6688* | Protein of unknown function; expression decreases by benomyl treatment or in an azole-resistant strain overexpressing MDR1 | -2.389 |

**Table S7.** Strains used in this study.

| Strain | Genotype | Source |
| --- | --- | --- |
| SC5314 | *C. albicans* reference strain | 1 |
| TACA12H10 -pLUX-1 | *C. albicans* his1Δ/Δ arg4Δ/Δ tac1Δ::ARG4/tac1Δ::HIS1 ura3Δ/Δ:URA3 | This study |
| TACA12H10 -pLUX-TAC1-1-1 | *C. albicans* his1Δ/Δ arg4Δ/Δ tac1Δ::ARG4/tac1Δ::HIS1 ura3Δ/TAC1Δ:URA3 | This study |
| UPC2M4A | *C. albicans* upc21Δ::FRT/ upc2-2Δ::FRT | 2 |
| UPC2M2A | *C. albicans* upc21Δ::FRT/ UPC2 | 2 |
| CAI4+pKE4-NRG1 | ura3Δ/Δ:URA3-PrTEF1-NRG1 | unpublished |

**Table S8.**
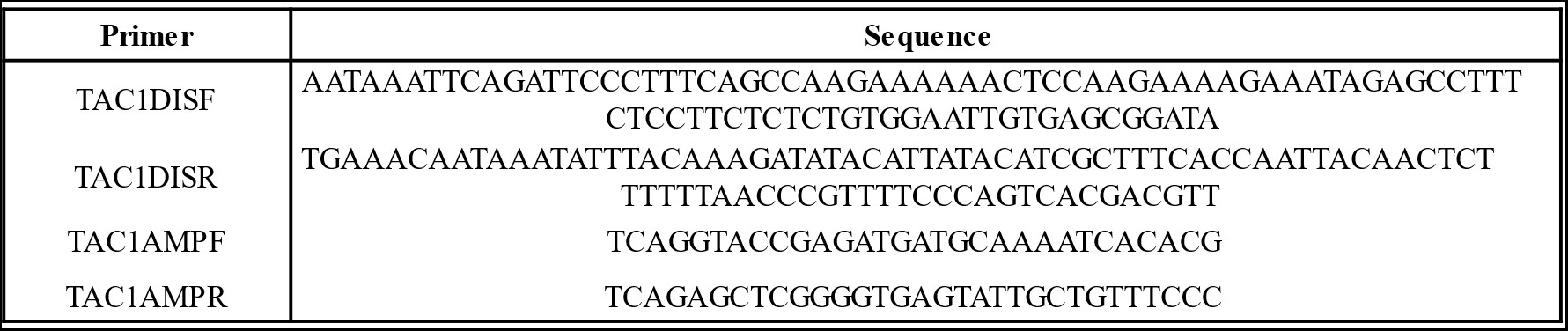
Oligonucleotides used in this study.

